# Midbrain organoids with an *SNCA* gene triplication model key features of synucleinopathy

**DOI:** 10.1101/2021.04.12.439480

**Authors:** Nguyen-Vi Mohamed, Julien Sirois, Janani Ramamurthy, Meghna Mathur, Paula Lépine, Eric Deneault, Gilles Maussion, Michael Nicouleau, Carol X.-Q. Chen, Narges Abdian, Vincent Soubannier, Eddie Cai, Harris Nami, Rhalena A. Thomas, Mahdieh Tabatabaei, Lenore K. Beitel, Karamjit Singh Dolt, Jason Karamchandani, Tilo Kunath, Edward A. Fon, Thomas M. Durcan

## Abstract

*SNCA,* the first gene associated with Parkinson’s disease, encodes the α-synuclein (α-syn) protein, the predominant component within pathological inclusions termed Lewy bodies (LBs). The presence of LBs is one of the classical hallmarks found in the brain of patients with Parkinson’s disease, and LBs have also been observed in patients with other synucleinopathies. However, the study of α-syn pathology in cells has relied largely on two-dimensional culture models, which typically lack the cellular diversity and complex spatial environment found in the brain. Here, to address this gap, we use 3D midbrain organoids (hMOs), differentiated from human induced pluripotent stem cells derived from patients carrying a triplication of the *SNCA* gene and from CRISPR/Cas9 corrected isogenic control iPSCs. These hMOs recapitulate key features of α-syn pathology observed in the brains of patients with synucleinopathies. In particular, we find that *SNCA* triplication hMOs express elevated levels of α-syn and exhibit an age-dependent increase in α-syn aggregation, manifested by the presence of both oligomeric and phosphorylated forms of α-syn. These phosphorylated α-syn aggregates were found in both neurons and glial cells and their time-dependent accumulation correlated with a selective reduction in dopaminergic neuron numbers. Thus, hMOs from patients carrying SNCA gene multiplication can reliably model key pathological features of Parkinson’s disease and provide a powerful system to study the pathogenesis of synucleinopathies.

## Introduction

Parkinson’s disease (PD) is the second most prevalent neurodegenerative disorder, affecting more than 10 million people worldwide ^1^. Genetic factors contribute to the complex pathogenesis of PD with approximately 5% of all patients suffering from a monogenic form of PD. *SNCA*, the first gene associated with familial PD, encodes the protein α-synuclein (α-syn) ^2^. Missense mutations in the *SNCA* gene cause a rare, autosomal dominant inherited form of PD ^3–8^. Moreover, copy number variations (CNVs) in the *SNCA* gene were also identified in patients with PD ^9, 10^. Indeed, the clinical phenotype of *SNCA* duplications resembles typical late-onset sporadic PD whereas *SNCA* triplications lead to a more widespread neurodegeneration with early-onset parkinsonism and dementia ^9, 11–14^. This implies that there is a direct relationship between *SNCA* gene dosage and disease severity ^9, 15^. Beyond these rare genetically determined forms, the abnormal accumulation of α-syn in the brain is a classical pathological hallmark for a group of related neurodegenerative disorders, collectively referred to as synucleinopathies. These include the vast majority of patients with sporadic PD, as well as those with Dementia with Lewy Bodies (DLB) and Multiple System Atrophy (MSA).

A cardinal feature of synucleinopathies is the pathological misfolding and aggregation of α-syn, and its accumulation within inclusion bodies in the brain, termed Lewy bodies (LBs), Lewy neurites (LNs) and Glia Cytoplasmic Inclusions (GCIs). A significant proportion of α-syn in these inclusions is phosphorylated at serine 129 (pS129Syn), a post-translational modification that has been detected in newly formed α-syn aggregates ^16^. The precursor to these phosphorylated aggregates is oligomeric α-syn, another important feature of α-syn pathology that is commonly observed in post-mortem PD brains. Oligomeric forms are believed to represent the initial seeds following misfolding of native α-syn, which can then induce further templating and aggregation of α-syn into phosphorylated aggregates ^17, 18^. These oligomers are believed to play a neurotoxic role by inducing mitochondrial and proteasomal defects, endoplasmic reticulum stress, inflammatory responses, and synaptic, autophagic and lysosomal dysfunction ^19^. Thus, while both oligomeric and phosphorylated α-syn aggregates have been shown to occur in synucleinopathies, how they form, and their precise effects have remained elusive.

To date, the majority of studies examining α-syn pathophysiology have relied on two-dimensional (2D) cell culture systems and rodent models ^20, 21^. With the advent of human induced pluripotent stem cells (iPSCs) and three-dimensional (3D) brain organoids, it is now possible to use patient-derived models to more faithfully reconstitute human brain-region specific features of the disease *in vitro*. One such region is the midbrain which contains the substantia nigra (SN), the area most affected in PD and where the majority of dopaminergic neurons (DNs) are lost. Single cell sequencing analysis of the SN demonstrated a high complexity of cell types, including neurons, astrocytes and oligodendrocytes and showed that the common genetic risk for PD is associated with DN-specific gene expression ^22^. Recent protocols to generate human midbrain organoids (hMOs) from iPSCs have made it possible to generate 3D human tissue *in vitro* that more closely resembles the native environment and cellular diversity, including DNs, found *in vivo* in the SN ^23^. Neurons within hMOs form interconnected networks, are organized in multiple layers, and exhibit a number of functional properties normally displayed by neurons, that include synapse formation and spontaneous electrophysiological activity ^24^. Thus, hMOs provides a human model in a dish with the potential to capture the 3D architecture, cellular diversity and connectivity found in the midbrain *in vivo*. In contrast to *in vivo* rodent models, human iPSC-derived organoids also have the advantage of being generated from patients and are thus more likely to faithfully recapitulate disease features^25-27^.

To investigate whether key features of synucleinopathies could be detected in a brain organoid model, we generated hMOs from an iPSC line derived from a patient with an *SNCA* gene triplication (*SNCA* Tri). These were compared to hMOs generated from isogenic control iPSCs derived from the same line, but in which the *SNCA* gene copy-number had been corrected to wild-type, through CRISPR/Cas9 genome editing. Using a combination of molecular, biochemical, immunocytochemical and flow cytometry approaches, we report an age-dependent increase in ɑ-syn levels and aggregation, including oligomeric- and S129-phosphorylated α-syn, in the *SNCA* Tri but not the control hMOs. Interestingly, pS129Syn was detected in both neurons and glial cells and correlated with a reduction in the number of DN over time, coinciding with broader neuronal loss and, conversely, an increase in the number of glial cells. Taken together, our findings support the idea that hMOs from patients with *SNCA* CNVs can model key pathological features of synucleinopathies and provide a novel platform to investigate the mechanism of α-syn pathogenesis in PD.

## Materials and methods

### Cell-line information and ethical approvals

The use of iPSCs in this research is approved by the McGill University Health Centre Research Ethics Board (DURCAN_IPSC / 2019-5374). SNCA lines (originally named AST23, AST23-2KO, AST23-4KO) were provided and generated by Dr Tilo Kunath from The University of Edinburgh according to methodology described in ^28^. The healthy individual line in **Figure S2** is named the NCRM-1 cell-line (NIH CRM, http://nimhstemcells.org/crm.html) while the SNCA Tri (b) line is named ND34391*H (Coriell, https://www.coriell.org/0/PDF/NINDS/ipsc/ND34391_CofA.pdf).

### Maintenance of iPSCs and hMO differentiation

For generation of hMOs, iPSCs were used only after a minimum of two passages following being thawed and were not passaged more than ten times. Details on iPSC passaging is found in our QC workflow for iPSC maintenance ^29^. hMOs were generated according to the published method ^30^. Briefly, 10,000 cells from a single cell suspension of iPSCs were seeded in each well of a 96 well ultra-low attachment plate, containing neuronal induction media. Plates were then centrifuged for 10 min at 1200 rpm to aggregate the cells. After 48 h, embryoid bodies (EBs) formed and the medium was replaced with fresh neuronal induction media. After 2 days, the medium was replaced with midbrain patterning medium, driving the EBs toward the midbrain fate. EBs were then embedded, 3 days later, in Matrigel® reduced growth factor and incubated for 24 h in tissue induction medium. Finally, the following day, embedded hMOs were transferred into a 6-well ultra-low attachment plate filled with final differentiation medium and cultured on an orbital shaker until required for experiments ^30^. For quality control purposes, mycoplasma tests were run monthly on iPSC cells and hMOs to ensure the absence of contamination.

### RNA isolation, cDNA synthesis, and qPCR

RNA was purified from iPSCs and hMOs using a NucleoSpin RNA kit (Takara) according to the manufacturer’s instructions. cDNA was generated using iScript Reverse Transcription Supermix (BioRad). Quantitative real-time PCR was performed on QuantStudio 3 Real-Time PCR System (Applied Biosystems) using the primers listed in **S2 Table**. This enabled a total reaction volume of 10 µl, including 5 µl of fast advanced master mix (Thermofisher Scientific), 1 µl 20X Taqman assay (IDT), 1 µl of cDNA and 3 µl of RNAse free water. Relative gene expression levels were analyzed using the Comparative CT Method (ΔΔCT method). The results were normalized to the GAPDH expression. The relative quantification (RQ) was estimated according the ΔΔCt methods^31^.

### Sanger sequencing of *SNCA* regions in iPSC lines

DNA was extracted from iPSC cultures using the Genomic DNA Mini Kit -Blood/Cultured Cell - (Genaid). The PCR reaction contained 1x Q5 Reaction Buffer (NEB), a detergent-free buffer containing 2.0 mM Mg^++^ at final 1x concentration, 200 µM dNTPs, 0.5 µM of each primer, 0.2U/µL of Q5 High-Fidelity DNA Polymerase (NEB) and 50-100 ng of DNA. Deletions were amplified from genomic DNA at 61°C and 63°C, while the *SNCA* coding region (without intron) was amplified at 65°C from cDNA. The amplification cycle was 2 min of denaturation at 96°C, followed by 35 cycles of 30 sec at 96°C, 30sec at 61°C for the 112bp deletion, 63°C for 231bp and 24bp deletions or 65°C for *SNCA* cDNA, then 1 min at 72°C, with a final extension of 2 min at 72°C. PCR products were purified using ExoSAP-IT PCR product cleanup reagent (ThermoFisher). DNA sequencing reactions were performed using the BigDye v3.1 cycle sequencing kit (ThermoFisher), using 0.3 µM of the corresponding single sequencing primer and 50 ng of purified DNA product. Sequencing reactions were placed on a thermocycler programmed for denaturation at 96°C for 2 min, followed by 25 cycles at 96°C for 20 sec, 50°C for 10 sec, and 60°C for 4 min. Sequencing reactions were purified using a BigDye XTerminator Purification kit (Thermo-Fisher). Sequencing was carried out on an Applied Biosystems SeqStudio Genetic Analyzer (ThermoFisher).

### Determination of *SNCA* copy number

The quantification of *SNCA* copy number was achieved using a combination of the Bio-Rad droplet digital PCR (ddPCR) QX200^TM^ system and a TaqMan^®^ assay which includes two sets of PCR primers and two DNA probes fused with different fluorophores (FAM and HEX). Locked Nucleic Acid (LNA^®^) DNA probes were used and designed according to manufacturer’s protocol (IDT). One primer/probe set was specific to the *SNCA* gene and the other primer/probe set was specific to an internal autosomal control gene *CDH2*. Near-confluent iPSCs cultured in a 6-well plate format were washed with PBS and treated with Accutase^®^ for 10 min at 37°C. Genomic DNA was extracted using the DNeasy extraction kit according to the manufacturer’s protocol (Qiagen). To prepare a typical ddPCR master mix for one 8-well strip, we mixed 90 µl of “ddPCR Supermix for Probes (no dUTP)” (Bio-rad), 75 µl H_2_O, 1.6 µl of each primer (stock 100 µM) and 0.5 µl of each probe (stock 100 µM). 19 µl of this master mix was distributed per well in an 8-well PCR tube strip. Two µl of the DNA extract was added into each corresponding well. The ddPCR droplet generation, reading, and quantification were performed according to the manufacturer’s protocol, and the ratio of *SNCA* alleles vs *CDH2* alleles calculated.

### Genomic stability testing

iPSC lines were sent to WiCell to be karyotyped according to WiCell’s sample preparation recommendations. For testing of hotspot mutations within the genome of the iPSCs used, genomic DNA was extracted with the Genomic DNA Mini Kit (Blood/Cultured Cell) (Geneaid). Genomic stability was then tested with the hPSC Genetic Analysis Kit (Stemcell, 07550) according to our earlier studies ^29^.

### Fixation and immunofluorescence staining of hMO cryosections

hMOs were fixed and cryosectioned according to our published methods ^30^. For staining, 14 μm cryosections were rehydrated in PBS for 15 min and surrounded with a hydrophobic barrier drawn with a barrier pen. The sections were then incubated with a blocking solution (5% normal donkey serum, 0.05% BSA, 0.2% Triton X-100 in PBS) for 1 h at RT in a humidified chamber. The sections were subsequently incubated overnight at 4°C with primary antibodies diluted in blocking solution (**Table S1**). The following day, cryosections were washed three times in PBS (15 min each) and then incubated in secondary antibodies diluted in a blocking solution for one hour at RT (**Table S1**). Sections were next washed three times in PBS for 15 min each, incubated with Hoechst (1/5000 in PBS) for 10 min, and finally washed once in PBS for 10 min. Finally, sections were mounted (Aqua-Poly/Mount, Polysciences), and the images acquired with a Leica TCS SP8 confocal microscope.

### Preparation of hMO cell lysates for immunoblotting

hMOs were washed with PBS and lysed through a syringe (BD #329420) in RIPA buffer [50 mM Tris HCl pH 7.5, 150 mM NaCl, 0.5% Triton X-100, 0.5% sodium deoxycholate (SDC), 1% sodium dodecyl sulfate (SDS), 1 mM dithiothreitol (DTT), 50 mM NaF, 5 mM Na3VO4] containing a protease inhibitor cocktail (Complete EDTA-free from Roche Diagnostics, Indianapolis, IN) and a phosphatase inhibitor cocktail (2×PhosSTOP from Roche Diagnostics). Proteins were quantified using the Bio-Rad DC Protein assay (Bio-Rad). The lysates were mixed with 4X Laemmli buffer and boiled for 5 min. Equal amounts of proteins were loaded for each sample and electrophoresed on a 10% polyacrylamide gel. Following SDS-PAGE separation, proteins were transferred onto nitrocellulose membranes. To increase the sensitivity of α-syn detection, membranes were fixed for 30 min at RT in 4% paraformaldehyde and 0.1% glutaraldehyde ^32^. Next, membranes were incubated in a 5% milk solution diluted in Tris-buffered saline containing 0.2% Tween^®^20 (TBST) for 1 hr. Membranes were incubated with primary antibodies overnight at 4°C (**Table S1**), then washed in 0.2% TBST and incubated with peroxidase-conjugated secondary antibodies for 1 h at RT. Membranes were again washed and revealed by chemiluminescence (Amersham Pharmacia Biotech, Quebec, Quebec, Canada). Image acquisition and densitometry were performed with a ChemiDoc™ MP System.

### Fontana Masson staining and colorimetric extraction

hMOs were fixed with 4% PFA for 1h at RT, then formalin fixed, dehydrated and paraffin infused using a tissue processor (HistoCore PEARL, Leica). The hMOs were then embedded in paraffin blocks before being sectioned using a microtome (RM2235, Leica). Fontana Masson staining was performed using an Abcam kit (#ab150669), following manufacturer’s instructions on 4 μm paraffin sections of hMOs. Entire hMOs were then imaged with the ZEISS Stemi 508 stereomicroscope combined with the ZEISS Axiocam ERc 5s camera. Images of sections were acquired with a clinical Olympus BX46 microscope and an Olympus DP27 digital color camera. Black dots from Fontana Masson stainings pictures were extracted using colorimetric selection from GIMP software (version 2.8.22) and quantified by ImageJ (version 2.0.0-rc-69/1.52i) following the method described ^33^.

### Synuclein proximity ligation assay (Syn-PLA) on paraffin sections

The study involving substantia nigra sections from human subjects was approved by the C-BIG Biorepository Review Ethics Board (REB #IRB00010120). Fixed hMOs were processed and paraffin embedded. Blocks were then cut into 4 μm sections using a microtome, dewaxed and rehydrated as described in Roberts et al. ^19^. Briefly, Syn-PLA involves four steps: antigen retrieval and recognition, ligation, amplification, and detection. For the antigen retrieval step, sections were microwaved in 10 mM of sodium citrate buffer pH6, washed in TBS with 0.1% Triton X-100, and blocked 1 h in 1 M glycine, 10% normal goat serum, and TBS with 0.1% Triton X-100. Sections were then incubated overnight at 4°C with antibodies targeting α-syn and pS129Syn (**Table S1**) for the antigen recognition step. The next day, after three washes in TBS with 0.1% Triton X-100, sections were incubated in secondary antibodies for 1h at RT and subsequently washed three times in TBS with 0.05% Tween^®^20.

For the ligation, PLA probes and diluent materials were obtained from Sigma-Aldrich kits (#DUO92009, #DUO92010 and #DUO92007). Sections were blocked in a PLA blocking solution, and then incubated in PLA probes combined with an ɑ-syn antibody overnight at 4°C according to ^34^ . After three washes, sections were incubated in a ligation solution for 1h at 37°C. Next, sections were incubated in amplification and detection solution for 2.5 h at 37°C. Sections were finally washed four times in TBS and stained with Hoechst (diluted 1/5000 in TBS) for 10 mins, before being mounted with an aqua-mounting media. Sections were then visualized under Leica TCS SP8 microscope, with consistent gain and laser settings across conditions.

### Immunofluorescence quantification of hMOs

The intensity of immunofluorescence signals and the area of staining for each antibody were quantified using an in-house developed Organoid Quantification (OrgQ) macro script (https://github.com/neuroeddu/OrgQ) that used the ParticleAnalyzer function within ImageJ. The script, written in Python, used greyscale channel images and the merged image taken from the confocal microscope as inputs. A mask outlining the organoid from the merged image was first created to account for the organoid area by using an automatically chosen threshold. This mask was next applied to channel inputs, so the quantification of the organoid area remained consistent within each channel. The script next loops through each image within a folder (each channel) for which a given threshold was chosen per image (or were pre-set from a csv file) to count all the particles based on the intensity. This analysis yielded the total number of particles for each channel quantified. At the same time, it also measured the area of each particle within each channel to quantify the sum of particulate areas for a specific channel, as well as to measure the total area of the organoid. All quantified data were outputted in a csv file.

### Flow cytometry analysis of dissociated hMOs

First, hMOs were harvested, washed with 1x D-PBS (MultiCell) and treated with TrypLE Express (Thermo-Fisher) three times (2x 10 min, 1x 5 min) at 37°C to create a single cell suspension. Between each incubation period, hMOs were triturated with 1 mL pipet tips. The TrypLE reaction was stopped by adding 1x D-PBS. The single cell suspension was next filtered through a 30 µm mesh (Miltenyi Biotec) and cells pelleted by centrifugation at 300g for 5 min. Pelleted cells were resuspended in 1x D-PBS and viability staining with Fixable Live/Dead Aqua (Invitrogen) was performed for 30 min at RT (protected from light). Cells were then washed with 1x D-PBS and centrifuged at 300g for 5 min. The cell pellet was next resuspended in FACS Buffer (5% FBS, 0.1% NaN3 in D-PBS) containing a Fc receptor blocking solution, Human TruStain FcX^TM^ (Biolegend). Protected from light, extracellular antigen stainings were performed. The optimal concentration (determined by titration) of extracellular antibodies (**Table S1**) was added to the cells suspended in FACS Buffer. These cells were incubated for 30 min at RT and protected from light. For intracellular staining (pS129Syn, α-syn, TH), the single cell suspension was fixed with a FIX & PERM Cell Fixation & Cell Permeabilization Kit (Thermo-Fisher) according to the manufacturer’s protocol. After fixation, cells were washed with FACS Buffer and centrifuged at 350g for 5 min. The optimal concentration (determined by titration) of pS129Syn antibody (**Table S1**) was added to the cells in PERM Buffer. The intracellular staining was performed for 20 min at RT and protected from light. The cells were washed twice with FACS Buffer and centrifuged at 350g for 5 min. After this step, cells were resuspended in FACS buffer and were then ready for flow cytometry analysis. For flow cytometry settings and parameters, voltages were set according to optimal PMT sensitivity using the peak 2 (Spherotech) voltration technique described previously by Maeker and Trotter ^35^. Compensation control was performed with Ultracomp beads (Thermo Fisher) using optimal antibody concentrations determined by titration. All data was acquired on the Attune NxT (Thermo-Fisher). Finally, all data generated was analyzed with FlowJo (Version 10.6) (Becton-Dickinson Biosciences). For pS129Syn and α-syn signals, after gating out debris and dead cells, we reduced the probability of false positives (i.e. type I errors) by gating three standard deviations above the mean, using the *SNCA* KO line as a negative control, in order to reduce the size of the population.

### Statistical analysis

Statistical analysis was performed using GraphPad Prism 8.4.1 software. All statistical tests were selected according to the normal distribution of data tested. Statistical significance was evaluated with a two-tailed unpaired t-test or a two-tailed Mann-Whitney test when two conditions were compared. A one-way analysis of variance (ANOVA) test followed by a Tukey’s multiple comparison test or a Kruskal-Wallis test followed by a Dunn’s multiple comparisons test was performed for more than two groups comparison. The statistical analyses were performed using Prism 6.0c software.

## Results

### Generation of midbrain organoids from patient-derived iPSCs with an *SNCA* **triplication**

As one of the genetic defects associated with the development of PD, the *SNCA* triplication promotes elevated expression of α-syn, ultimately resulting in an increase in α-syn aggregation and an accompanying loss of DNs. To determine whether we could replicate the effects of this CNV event in a human model in a dish, in this study we differentiated hMOs from iPSCs reprogrammed from a patient carrying a triplication in the *SNCA* gene (*SNCA* Tri). As our control, we differentiated hMOs from iPSCs derived from the same patient line, but in which the *SNCA* allele copy number had been corrected to a wild-type state by deleting two of the four copies of the *SNCA* gene (Isog Ctl), using CRISPR/Cas9 genome editing. As an additional control, we differentiated hMOs from iPSCs derived from the same patient line in which all four *SNCA* alleles had been deleted (*SNCA* KO). To assess the quality of our three iPSC lines, we stained them for cell surface and pluripotency markers. All colonies expressed high levels of the glycoprotein antigen TRA1-60R, the glycolipid antigen SSEA-4, the transcription factor OCT3-4, and the transcription factor NANOG. Pluripotency markers were homogeneously observed in nearly all cells, which reflects the undifferentiated state of the iPSC cultures used as starting material for hMO generation (**Fig. S1A, Table S1**). Karyotype analysis showed no clonal abnormalities for any of the three lines (**Fig. S1B**). Genomic stability was assessed by monitoring copy numbers in critical hotspot regions that are often commonly mutated during reprogramming or extensive cell passaging ^29^. In these regions, we did not observe any of the recurrent abnormalities that have been reported ^36–38^ (**Fig. S1C**). At the DNA level, we also confirmed the number of *SNCA* alleles in each cell-line through digital droplet PCR (**Fig. S1D, Table S2**). As expected, we detected twice the copy number of the *SNCA* alleles relative to a control gene (CDH2) in *SNCA* Tri, compared to the Isog Ctl iPSCs, with no copies of the *SNCA* alleles in the *SNCA* KO iPSCs. Sequencing of the Isog Ctl line identified the break points of the three CRISPR/Cas9-induced deletions within the *SNCA* locus (**Fig. S1E, Table S2**). As only two of these are predicted to disrupt the α-syn protein coding sequence (the third being a small 24 bp deletion within an intron), the findings confirm the desired correction from four to two functional copies of the *SNCA* gene in the Isog Ctl line.

Following our previously published methods ^30^, we derived hMOs from the *SNCA* iPSC lines using classical midbrain patterning factors ^26, 39–45^ (**Fig. 1A**). Across batches, we observed that *SNCA* Tri hMOs were slightly smaller than Isog Ctl at 60 days (2.8±0.06 mm versus 3.1±0.06 mm mean diameter) (**Fig. 1B)**. This difference was more pronounced at 100 days (2.4±0.03 mm versus 3.1±0.06 mm mean diameter) (**Fig. 1B**). It has been reported that, upon exogenous treatment with dopamine, hMOs accumulate neuromelanin granules, a by-product of dopamine synthesis ^26, 30, 46^. Thus, we treated 35-day old hMOs for 10 days with 100 μm of dopamine followed by Fontana Masson staining of hMOs sections. We observed a similar increase in neuromelanin granules in *SNCA* Tri and Isog Ctl hMOs treated with 100 μm dopamine (**Fig. 1C**). Interestingly, we also noted spontaneous (without exogenous dopamine treatment) neuromelanin granule accumulation in long-term 6-month old hMOs cultures, as illustrated in **Fig. S1F**.

**Figure 1.**
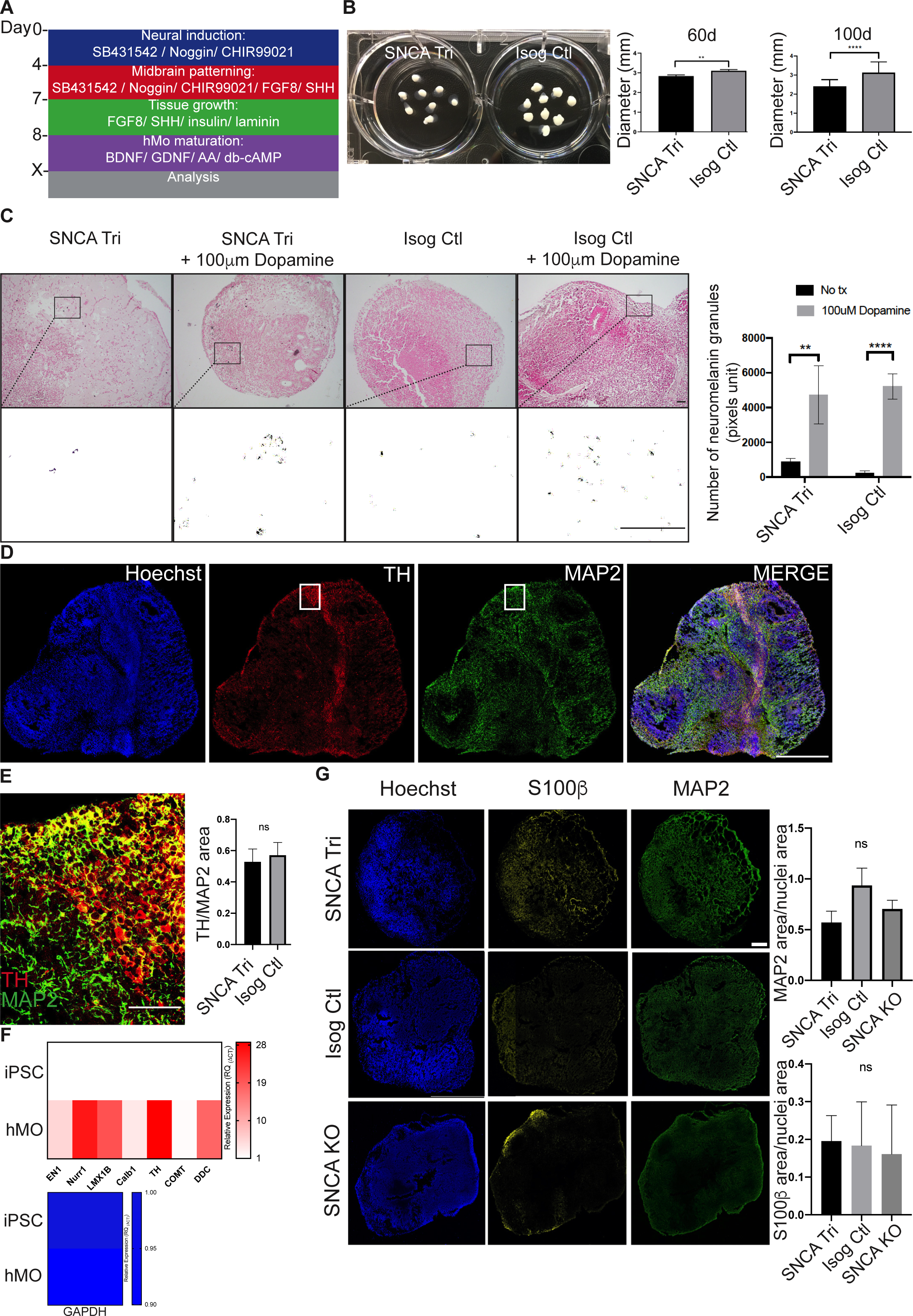
Generation of hMOs from *SNCA* lines. (**A**) Summary of patterning factors used to differentiate hMOs from iPSC lines. (**B**) *SNCA* Tri hMOs were smaller than Isog Ctl hMOs at 60 days (n=59, mean+/-SEM, unpaired t-test, two-tailed) and at 100 days (n=8, mean+/-SEM, unpaired t-test, two-tailed). (**C**) Fontana Masson staining showed that 45-day old hMOs treated for 10 days with 100 μm dopamine accumulated neuromelanin granules (n=6, mean+/-SEM, one-way ANOVA, Kruskal-Wallis test followed by Dunn’s multiple comparisons test). Scale bars = 200 μm. Black and white images correspond to a close-up of the neuromelanin pigments extracted from the pictures above. (**D**) Cryosections of 45-day old *SNCA* Tri hMOs stained for neurons (MAP2), dopaminergic neurons (TH) and nuclei (Hoechst). Scale bar = 1 mm. (**E**) Higher magnification of *(**D**)* white square (merged) showing neurons (MAP2 in green) and dopaminergic neurons (TH in red). Scale bar = 250 μm (graph: n=19, mean+/-SEM, unpaired t-test, two-tailed). (**F**) Real-time PCR depicting normalized expression level of midbrain RNA levels for EN1, Nurr1, LMX1B, Calb1, TH, COMT, DDC at 50 days in Isog Ctl hMOs, normalized to endogenous GAPDH and actin controls (n=3, mean+/-SEM). (**G**) Cryosections of 50 days old hMOs stained for neurons (MAP2), astrocytes (S100β) and nuclei (Hoechst). Scale bar = 250 μm. Quantification of MAP2 and S100β staining normalized to nuclei staining (n=6, mean+/-SEM, one-way ANOVA, followed by Tukey’s multiple comparisons test).

To further confirm the midbrain identity of the hMOs, we performed immunoblotting with 30-day old hMO lysates to detect the expression of common midbrain markers. Tyrosine hydroxylase (TH) is the rate-limiting enzyme in dopamine biosynthesis and a marker of catecholaminergic neurons. Immunoblots revealed lower TH protein levels in *SNCA* Tri compared to Isog Ctl hMOs lysates whereas there were no significant differences in the levels of TUJ1, a pan-neuronal marker (**Fig. S1G**). To further confirm that the organoids contained DNs, we stained cryosections from 50-day old hMOs for TH (**Fig. 1D**) as well as MAP2, a neuronal marker. Consistent with other published findings ^26, 27, 30^, we could detect the presence of DNs stained for TH within the hMOs (**Fig. 1D, E**). Quantification showed that the surface area of TH staining represented approximately 45% of the total MAP2 staining area, with no significant differences between Isog Ctl and *SNCA* Tri hMOs (**Fig. 1E**). In addition to immunocytochemistry, we used quantitative PCR (qPCR), to detect several transcripts known to be expressed in midbrain DNs that included engrailed homeobox 1 (EN1), nuclear receptor related 1 protein (Nurr1), LIM homeobox transcription factor 1-beta (LMX1B), calbindin 1 (Calb1), TH, catechol-O-methyltransferase (COMT) and dopa decarboxylase (DDC) in 50-day hMOs. As expected, their expression was absent in the parental iPSCs, whereas transcript levels of the housekeeping gene GAPDH were similar in both iPSCs and hMOs (**Fig. 1F)**. In addition to DNs, other cell types are also present within the hMOs and to assess the diversity of the cellular populations, we stained for MAP2, found primarily in neurons, and S100β, a marker for astrocytes (**Fig. 1G**). We observed similar staining for both cell types in sections from hMOs across all three genotypes. (**Fig. 1G**). The diversity of cell populations present in these hMOs was also confirmed in our earlier study by single cell RNA sequencing ^30^. Taken together, our findings indicate that hMOs from our patient and control lines display neurochemical, gene and protein expression, and cellular characteristics consistent with a midbrain identity.

### Elevated expression of α-syn in *SNCA* triplication midbrain organoids

Next, we sought to determine whether the increase in *SNCA* copy number in the patient iPSC line led to elevated synuclein transcription and translation. First, hMOs were cultured for 50 days, and total RNA purified from the hMOs, to quantify *SNCA* transcript levels by qPCR. We observed an approximate 2.5-fold increase of *SNCA* mRNA levels in *SNCA* Tri relative to Isog Ctl hMOs, whereas α-syn RNA levels were undetectable in *SNCA* KO hMOs (**Fig. 2A**, **Table S2**). Consistent with qPCR findings, α-syn protein levels, as measured by immunoblotting, were significantly elevated in the *SNCA* tri versus Isog Ctl hMOs, whereas α-syn was undetectable in *SNCA* KO hMOs (**Fig. 2B**). Staining for α -syn protein in cryosections of the 50-day old hMOs also revealed a higher expression in *SNCA* Tri compared to Isog Ctl hMOs (≃2 fold change), whereas α-syn staining was negligible in *SNCA* KO hMOs (**Fig. 2C, D**).

**Figure 2.**
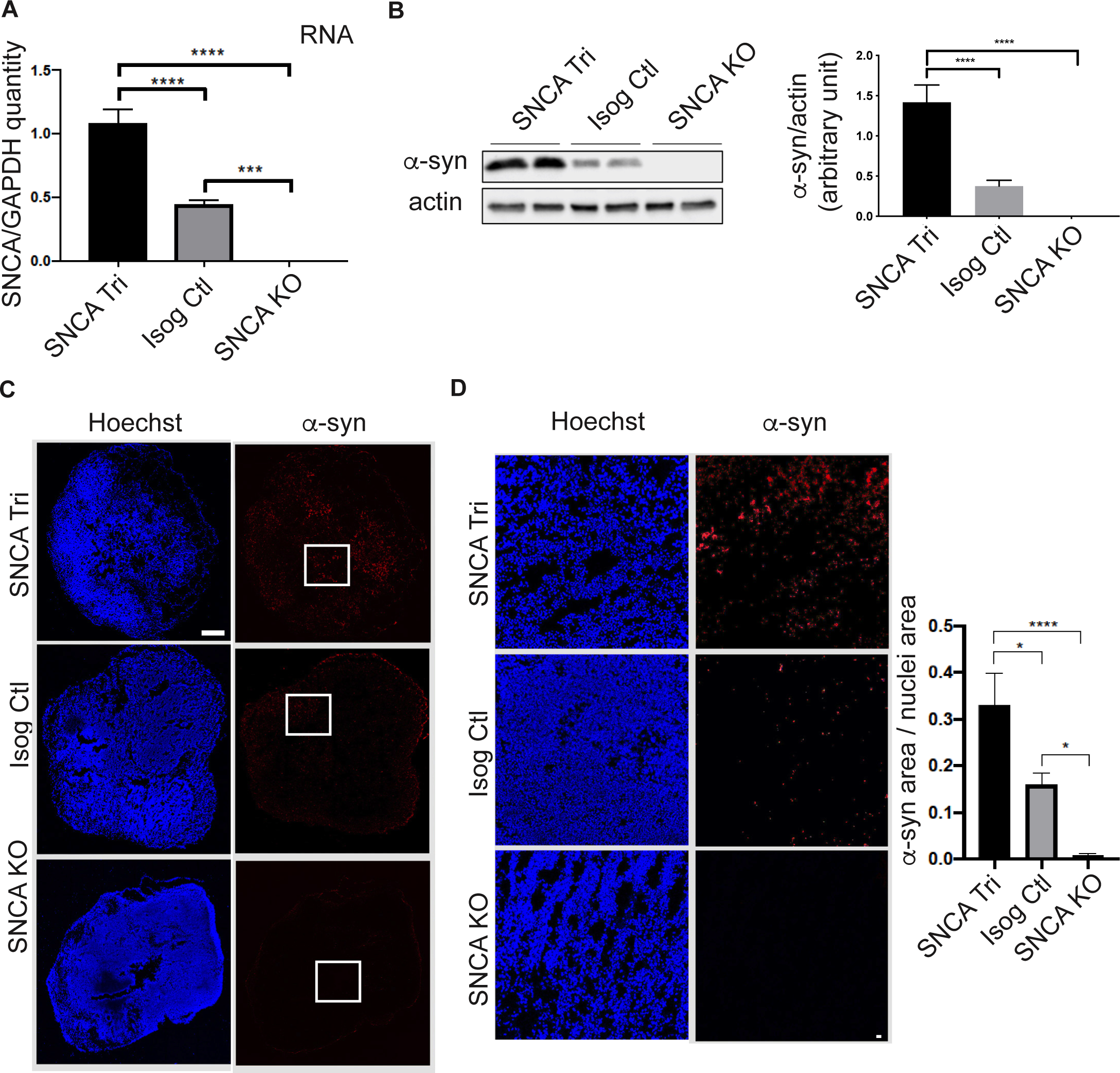
Increased *SNCA* mRNA and ɑ-syn protein level in *SNCA* Tri hMOs. (**A**) Real-time PCR depicting normalized expression level of *SNCA* RNA level at 50 days compared to endogenous GAPDH control (n=8, mean+/-SEM, one-way ANOVA, followed by Tukey’s multiple comparison test). (**B**) Western blot analysis of α-syn normalized to actin in 50-day old hMOs. Quantification by densitometry (arbitrary units) (n=8, mean+/-SEM, one-way ANOVA, followed by Tukey’s multiple comparisons test). (**C**) Cryosections of 50-day old hMOs stained for α-syn (α-syn), and nuclei (Hoechst). Scale bar = 250 μm. (**D**) Higher α-syn (α-syn) magnification of white squares in *(**C**)*. Quantification of α-syn staining normalized to nuclei staining (n=6, mean+/-SEM, one- way ANOVA, followed by Tukey’s multiple comparisons test). Scale bar = 50 μm.

We next extended the maturation time of the hMOs and repeated our analysis with 100-day old hMOs. After 100 days in culture, immunoblot analysis using two antibodies recognizing distinct epitopes demonstrated that α-syn was increased in the *SNCA* Tri versus Isog Ctl hMOs (**Fig. 3A**), consistent with findings at 50 days (**Fig. 2**). No signal was detected in the *SNCA* KO, further confirming the specificity of the α-syn signal. Staining of cryosections from the 100-day old hMOs also confirmed that α-syn levels were elevated in the *SNCA* Tri relative to Isog Ctl hMOs, with no discernible signal in *SNCA* KO sections (**Fig. 3B**). While the majority of α-syn signal was colocalized with MAP2^+^ (**arrows, Fig. 3C**), colocalization of α-syn was also observed in S100β^+^ astrocytes (**triangles, Fig. 3C**). Interestingly, in sections from *SNCA* Tri hMOs, we observed an intense α-syn signal in astrocytes and neurons (**stars, Fig. 3C**), indicative of inclusions or aggregates that could potentially be forming in both cell types. Overall, our findings demonstrate an increase in the expression of *SNCA* mRNA and α-syn protein in hMOs from the *SNCA* triplication line relative to the Isog Ctl and SNCA KO, detectable at 50 days and persisting in 100-day old hMOs.

**Figure 3.**
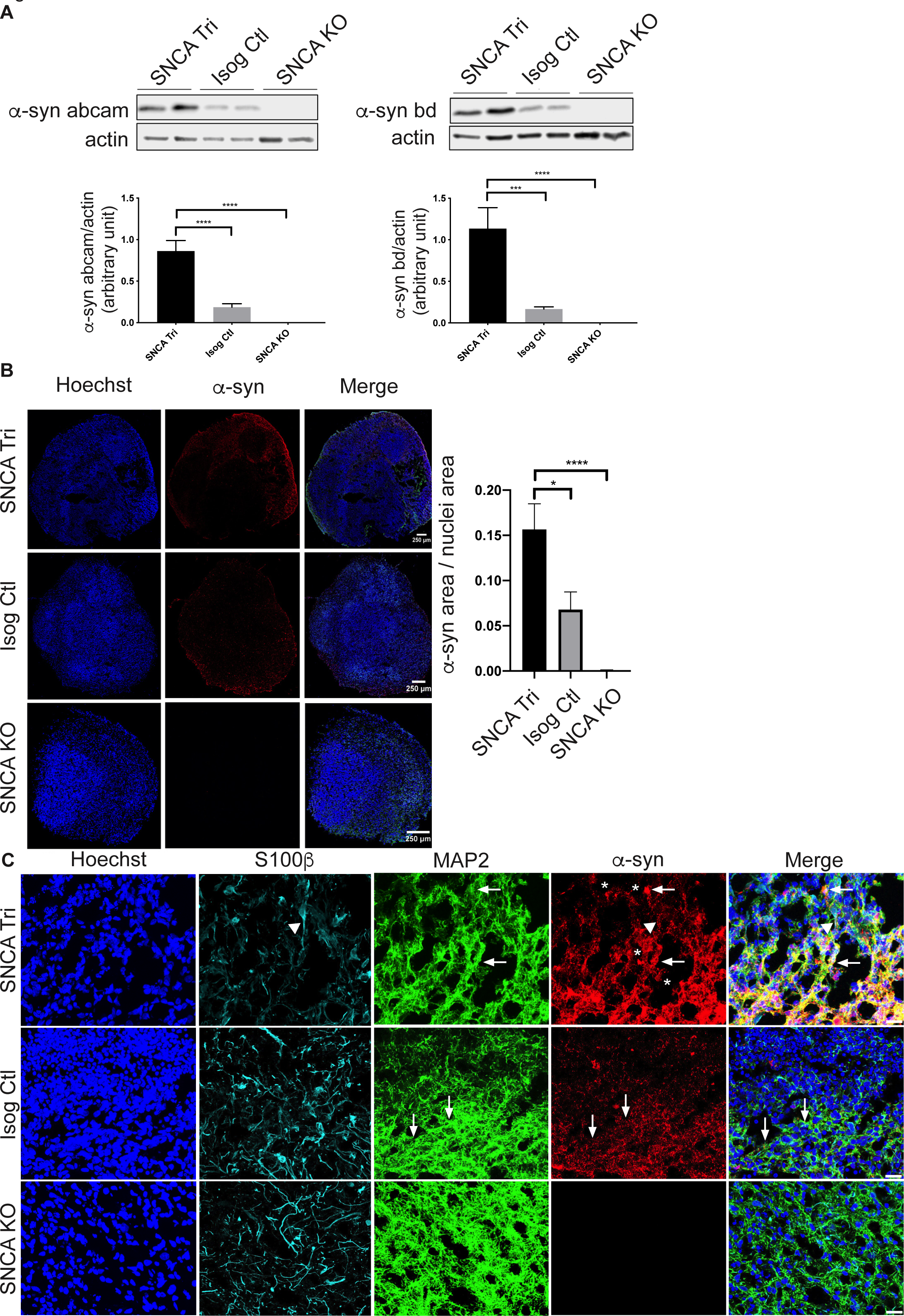
Increased ɑ-syn protein level in 100 days old hMOs. (**A**) Western blot analysis of α-syn normalized to actin in 100-day old hMOs. Quantification by densitometry (arbitrary units) (n=8, mean+/-SEM, one-way ANOVA, followed by Tukey’s multiple comparisons test). (**B**) Cryosections of 100-day old hMOs stained for α-syn (α-syn) and nuclei (Hoechst). Quantification of ɑ-syn staining normalized to nuclei staining (n=7, mean+/-SEM, one-way ANOVA, followed by Tukey’s multiple comparisons test). Scale bar = 250 μm. (**C**) Cryosections of 100-day old hMOs stained for α-syn (α-syn), neurons (MAP2), astrocytes (S100β), and nuclei (Hoechst). Scale bar = 100 μm.

### Oligomeric and phosphorylated ɑ-syn aggregates form over time in *SNCA* **triplication midbrain organoids**

PD is associated not only with elevated α-syn levels but also with its misfolding and aggregation. Both oligomeric and phosphorylated α-syn aggregates have been reported to be present in LBs during advanced stages of PD ^19^, leading to neuronal toxicity ^47, 48^. First, to detect misfolded and oligomeric forms of α-syn, we used an α-syn proximity ligation assay (Syn-PLA) ^34^ in which a signal is generated when two complementary PLA probes, recognizing interacting α-syn molecules (i.e., oligomers), bind in close proximity. The signal detected from each pair of PLA probes is then amplified and visualized as an individual fluorescent spot using confocal microscopy. This approach was first validated in post-mortem SN sections from a patient with DLB. We observed that the Syn-PLA signal colocalizes with α-syn and pS129Syn in cytoplasmic spherical LBs (**triangles, Fig. 4A**), as well as Lewy neurite structures (**arrows, Fig. 4A**). Similarly, we observed a robust accumulation in the oligomeric forms of α-syn in 100-day old *SNCA* Tri compared to Isog Ctl hMOs (**Fig. 4B, C**), and only a very faint background signal in *SNCA* KO hMOs. Consistent with this finding, we observed a similar pattern for Syn-PLA staining in hMOs derived from the *SNCA* triplication line, reprogrammed from a different patient with a SNCA triplication, compared to hMOs derived from a healthy individual with normal SNCA copy number (**Fig. S2A**). Interestingly, the Syn-PLA signal was consistently stronger than the pS129Syn signal in *SNCA* Tri hMOs, suggesting that this approach is sensitive for detecting α-syn oligomers and that they are an early manifestation of α-syn pathology, prior to the appearance of abundant pS129Syn-positive inclusions (**Fig. 4B**). Indeed, whereas pS129Syn was clearly increased in 100-day old *SNCA* tri hMOs lysates by immunoblotting (**Fig. 5A**), in cryosections, only a small proportion of cells within these hMOs were pS129Syn positive (**Fig. 5B**). Consistent with this finding, using paraffin-embedded sections, we observed only small, distinct populations of cells that were positive for pS129Syn staining in the hMOs derived from the second *SNCA* triplication line, compared to hMOs derived from our non-PD control line (**Fig. S2B, C**). Whereas pS129Syn occasionally colocalized with GFAP^+^ astrocytes (**triangle, Fig. 5C**), the majority of pS129Syn colocalized with neurons (MAP2^+^GFAP^-^) (**arrows, Fig. 5C**). Thus, *SNCA* Tri hMOs display features of α-syn pathology, including both oligomeric and phosphorylated forms of α-syn.

**Figure 4.**
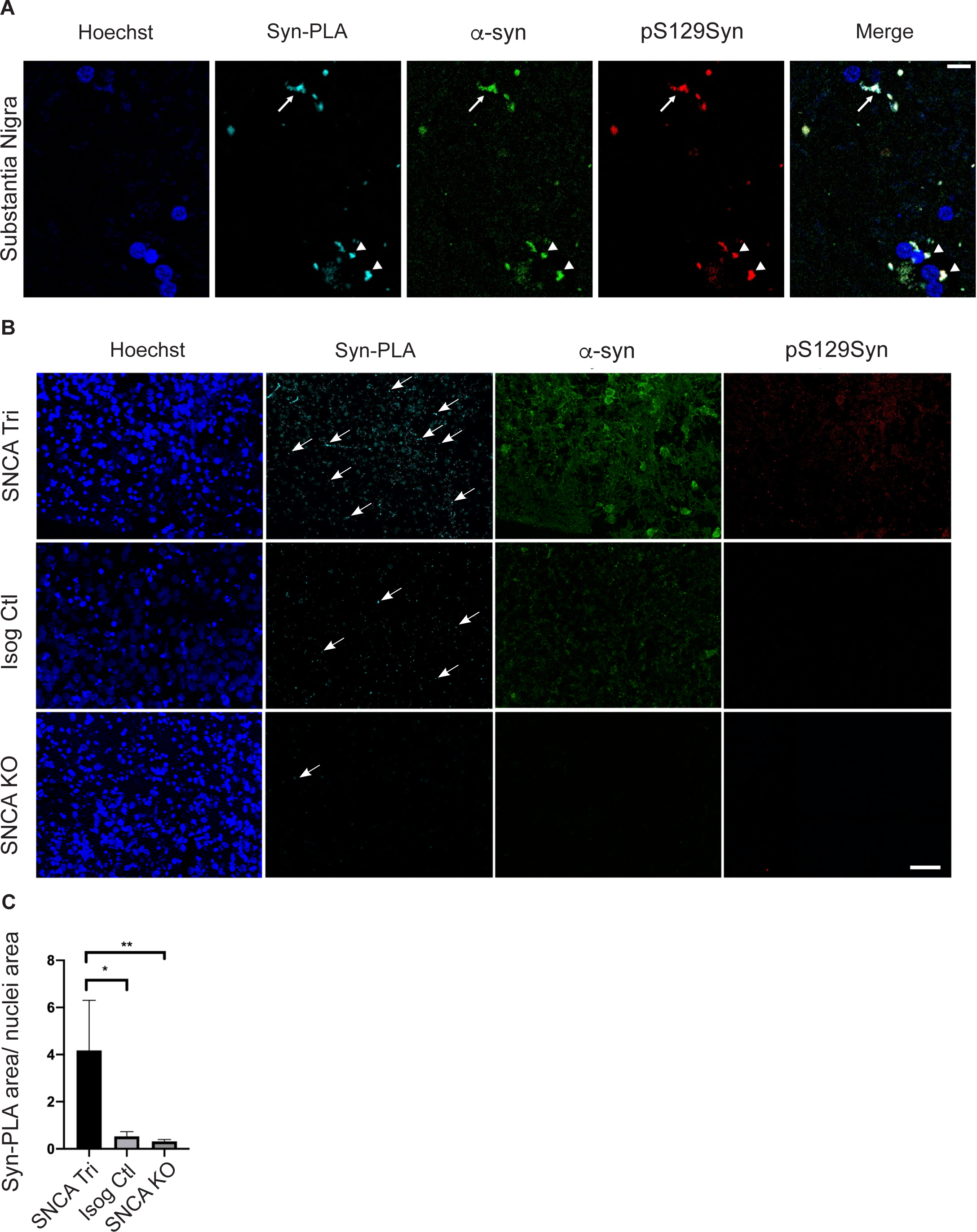
Oligomeric ɑ-syn forms detected in 100-day old hMOs. (**A**) Syn-PLA on positive control paraffin sections, SN of patient with DLB, showing oligomeric α -syn forms (Syn-PLA), α-syn (α-syn), and pS129Syn in LNs (arrow) and LBs (triangles). (**B**) Syn-PLA on paraffin sections showed the highest quantity of α-syn oligomers (Syn-PLA), α-syn (ɑ-syn) and pS129Syn signal in *SNCA* Tri hMOs. Scale bar = 40 μm. (**C**) Quantification of Syn-PLA signal normalized to nuclei staining (n=9, mean+/-SEM, Kruskal-Wallis test, followed by Dunn’s multiple comparisons test).

**Figure 5.**
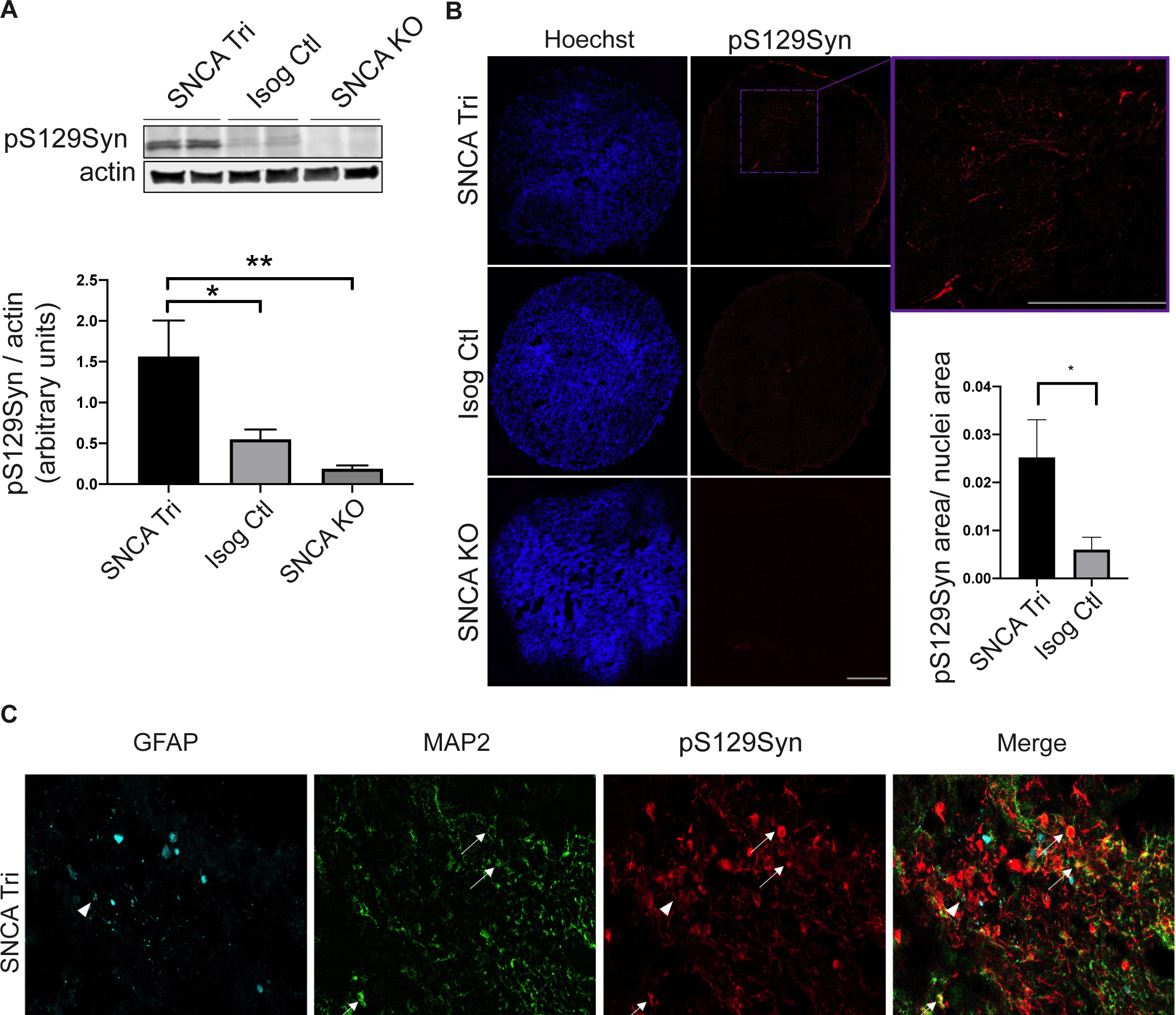
Endogenous pS129Syn observed in 100-day old hMOs. (**A**) Western blot analysis of pS129Syn normalized to actin in 100-day old hMOs. Quantification by densitometry (arbitrary units) (n=16, mean+/-SEM, one-way ANOVA, followed by Tukey’s multiple comparisons test). (**B**) Cryosections of 100-day old hMOs stained for pS129Syn and nuclei (Hoechst). Quantification of pS129Syn staining normalized to nuclei staining (n=8, mean+/-SEM, unpaired t-test, two-tailed). Scale bar = 250 μm. (**C**) Cryosections of 100-day old hMOs stained for pS129Syn, neurons (MAP2) and astrocytes (GFAP). Scale bar = 50 μm.

### Levels of phosphorylated α-syn aggregates increase with maturation of the hMOs

As hMOs are composed of multiple cell types ^23, 26, 27, 49^, they offer a unique model to explore the contribution of different cell populations on the ability of α-syn to aggregate. Our earlier findings with PLA and imaging of hMO cryosections showed aggregates were detected in both neurons and glia, albeit to varying degrees. To extend these findings further, we next performed single cell flow cytometry analysis. After dissociation of 100-day old hMOs into single cell suspensions, we performed cell surface staining for neuronal and glial cells as well as internal staining for α-syn and pS129Syn, to both quantify the levels of phosphorylated α-syn aggregates, and to measure the levels of these aggregates within both neuronal and glial cells (**Fig. 6A, Table S1**). Consistent with the results depicted in **Fig. 5**, we observed an enrichment of pS129Syn levels in the 100-day old *SNCA* tri hMOs (≃ 4 fold more in viable dissociated *SNCA* Tri cells) compared to the Isog Ctl hMOs, with cells from *SNCA* KO hMOs used as our negative control in the gating strategy (**Fig. 6B, C**). This was in the same range as the ≃4.2-fold increase in staining previously observed in the cryosections (**Fig. 5B**). Similar to the cryosections, pS129Syn signal was detected in less than 1% of dissociated *SNCA* Tri cells after 100 days (**Fig. 6C**). To investigate whether α-syn and pS129Syn accumulates further over time, the same analysis was performed on hMOs that were matured for 170 days in culture (**Fig. 6D-F Table S1**). Consistent with findings by immunoblotting and immunostaining at earlier time points (**Figs. 2 and 3**), we observed a significantly higher proportion of viable cells dissociated from *SNCA* Tri hMOs that express α-syn (≃11.1%) compared to those dissociated from Isog Ctl hMOs (≃5.5%) (**Fig. 6D histogram, S3A, S1 Table**). As expected from previous results, we also found a significant difference in the percentage of cells expressing pS129Syn in *SNCA* Tri hMOs relative to the Isog Ctl hMOs (≃0.74% vs ≃0.5%). Remarkably, we still observed that less than 1% of *SNCA* Tri cells were positive for pS129Syn (**Fig. 6E**), however, the amount of pS129Syn detected was almost double compared to the 100-day time point (≃0.74% versus ≃0.43%). As an additional control, we also verified that all cells which were pS129Syn positive were also α-syn positive in both genetic backgrounds (**Fig. 6F**).

**Figure 6.**
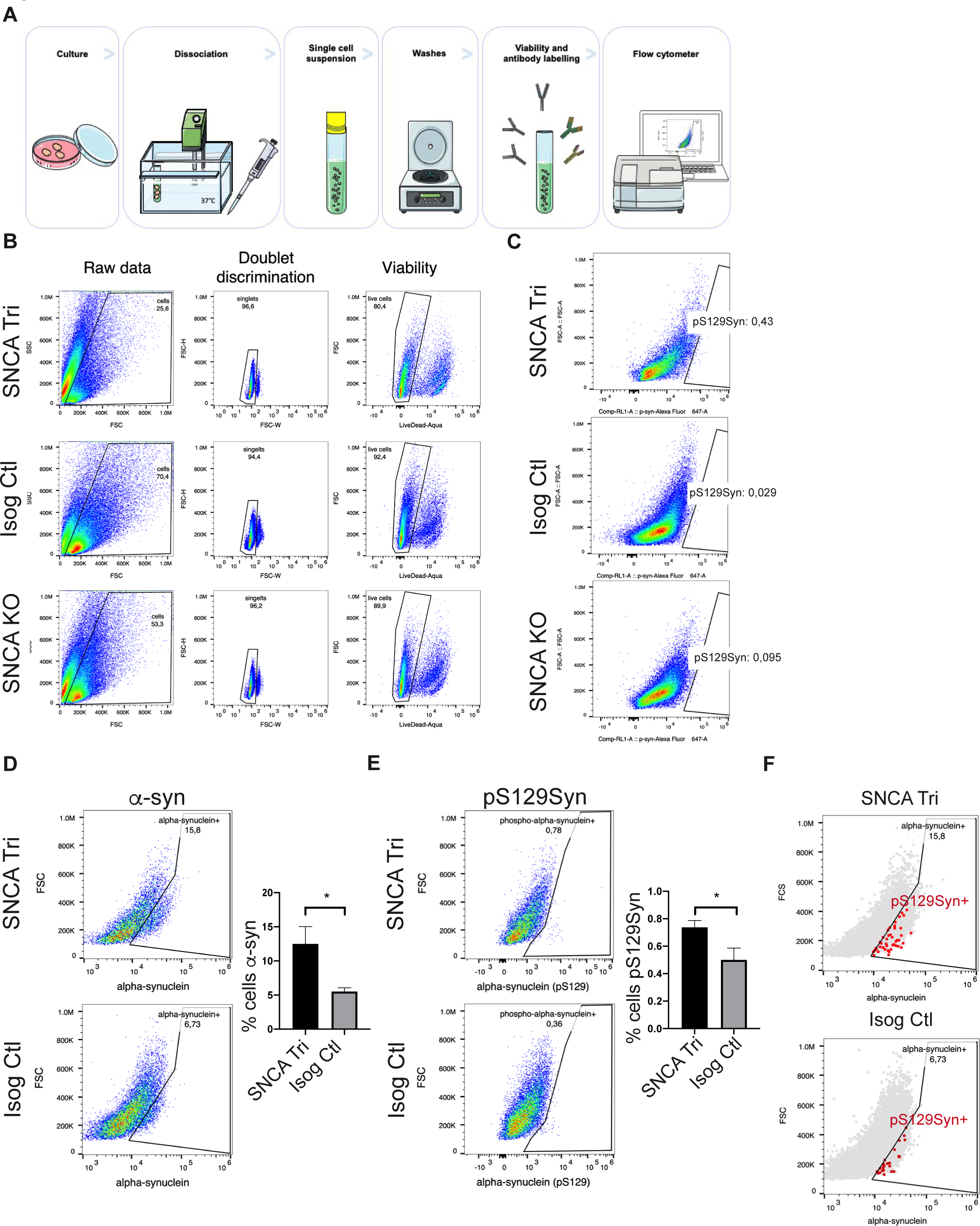
pS129Syn accumulation in *SNCA* Tri hMOs. (**A**) Flow cytometry sample processing schematic. Three organoids per genotype were incubated into enzymatic solution at 37°C, then manually dissociated with a pipet. Single cell suspensions were obtained after the filtration of dissociated tissues. The single cell suspensions were labelled for cell viability and antibodies against internal and external epitopes, before signals were read by the flow cytometer. (**B**) Gating strategy for the removal of cellular debris, doublet discrimination and cell viability. (**C**) Proportion of total cells carrying pS129Syn in single cell dissociated hMOs (3 hMOs pooled per line) measured by flow cytometry. (**D**) Quantification of percentage of cells carrying α-syn in dissociated 170-day old hMOs (n=5, 3 hMOs pooled per n, 15 organoids total, mean+/- SEM, unpaired t-test, two-tailed). The histogram represents 5n while the dot-plot represents 1n representative. (**E**) Quantification of percentage of cells carrying pS129Syn in dissociated 170-day old hMOs (n=6, 3 hMOs pooled per n, 18 organoids in total, unpaired t-test, two-tailed). (**F**) Control showing that all pS129Syn cells were also α-syn positive.

Findings from immunohistochemistry imply that the pS129Syn signal mostly colocalizes in neurons relative to glial cells. Thus, we next investigated in which cell-type the pS129Syn signal was present after 100 days in culture, a time point when pS129 was just beginning to accumulate. We used a combination of surface markers ^50, 51^ that included CD56 (NCAM-1, a neural cell adhesion molecule) and CD24 (a heat stable antigen receptor, nectadrin). Based on the literature, neurons were classified as CD56^+^CD24^+^ / CD56^+^CD24^-^/ CD56^-^CD24^+^ and glial cells as CD56^-^CD24^-^ ^50, 51^. The breakdown of the specific marker combinations is indicated in **Fig. S3B**. Quantification showed that, 75% of the pS129Syn signal was localized in neurons identified as CD56^+^CD24^+^, CD56^+^CD24^-^ and CD56^-^CD24^+^ whereas 25% was localized in CD56^-^CD24^-^ glia. For simplicity, in **Fig. 7A** we pooled and collectively depicted the neuronal populations as CD56^+^CD24^+^ and the glial population as CD56^-^CD24^-^. In *SNCA* Tri hMOs (green), we observed more pS129Syn in neurons than glial cells, whilst its presence in both cell populations in Isog Ctl was comparable to the background signal observed in *SNCA* KO hMOs (red and blue). Overall, our findings revealed that phosphorylated α-syn aggregates appear in both neurons and glial cells, albeit to varying degrees, with the levels increasing with maturation of the hMOs.

**Figure 7.**
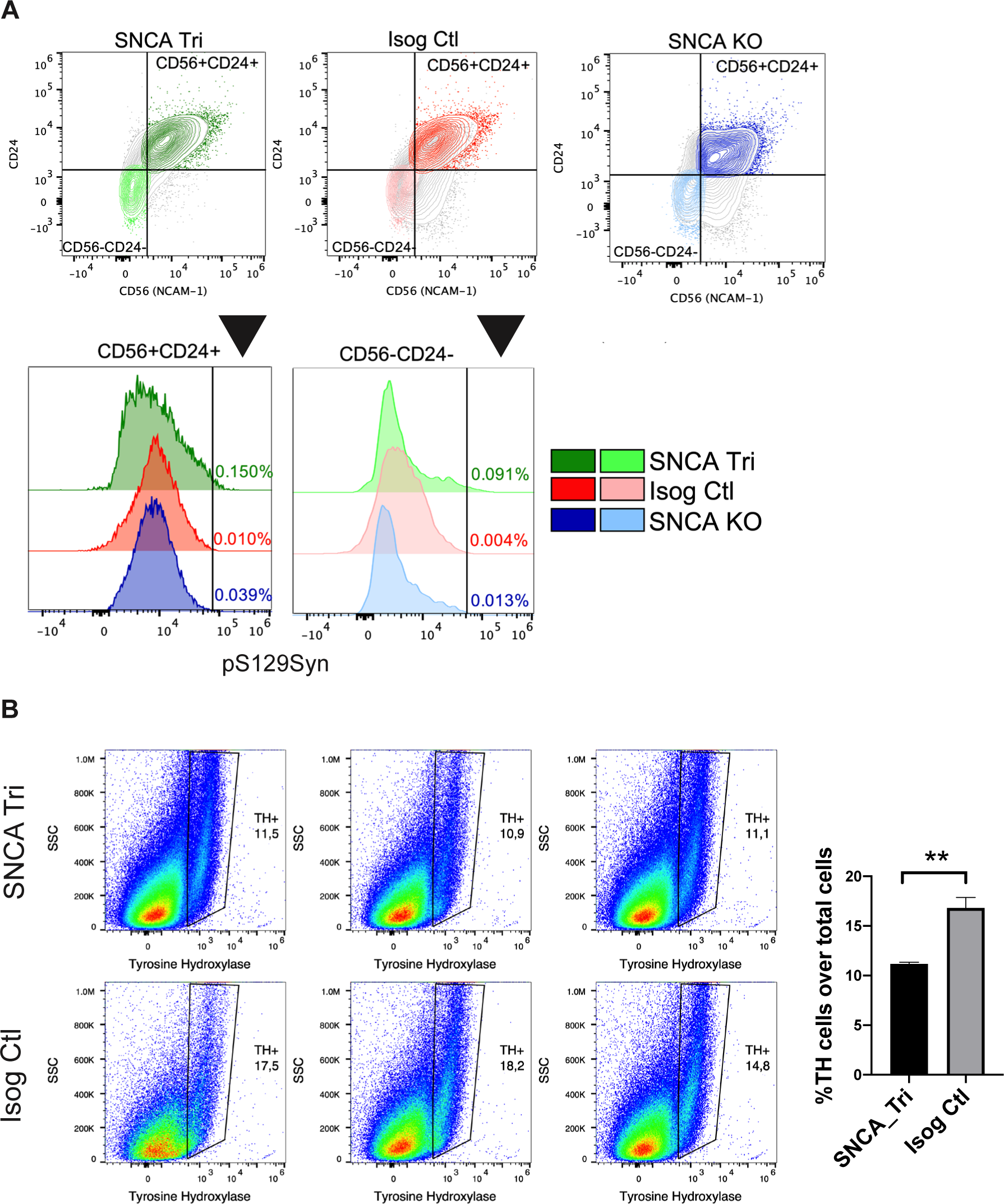
Localization of pS129Syn and dopaminergic neuron loss. (**A**) Proportion of CD56^+^CD24^+^neurons and CD56^-^CD24^-^ glial cells carrying pS129Syn (3 hMOs pooled per line) as measured by flow cytometry. (**B**) Quantification by flow cytometry of percentage of DN (TH-positive cells) in 100-day old hMOs (n=3, 3 hMOs pooled per n, 9 organoids total, mean+/-SEM, unpaired t-test, two-tailed).

### An increase in α-syn aggregates correlates with a reduction in dopaminergic neuron numbers

As synucleinopathies are characterized by the degeneration of vulnerable cell populations in the brain, we next investigated whether we could detect cell loss in hMOs and whether this correlated with the appearance of pS129Syn aggregates. Using a similar strategy as above, we quantified the levels of neuron (CD56^+^CD24^+^, CD56^+^CD24^-^, CD56^-^ CD24^+^) and glia populations (CD56^-^CD24^-^) by flow cytometry in 100-day hMOs. We detected similar proportion of neurons and glia in *SNCA* Tri and Isog Ctl hMOs (**Fig. S3C, D**). Using CD49f^+^ as an astrocytes marker ^52^, we noted a tendency for astrocyte enrichment in *SNCA* Tri hMOs, but this did not reach significance in our analysis (**Fig. S3E**). In contrast, after 170 days in culture, we observed a significant decrease in the proportion of neurons and an increase in glial cells in *SNCA* Tri hMOs compared to Isog Ctl **(Fig. S3F)**, implying that as neuronal numbers start decreasing, increased levels of gliosis may be occurring. This was further supported by our observations of a significant reduction in the number of TH-positive neurons in 100-day old *SNCA* Tri hMOs compared to Isog Ctl (**Fig. 7B**). We must however point out that there was no decrease in the overall number of total neurons at 100 days (**Fig. S3C**) which may suggest that DN are selectively vulnerable, and when lost are replaced by other neuronal subtypes.

Taken together, hMOs with a PD-associated triplication in the *SNCA* gene display increased levels of α-syn and α-syn aggregation, coinciding with a reduction in TH-positive neurons over time. Overall, our findings demonstrate that hMO can model key pathological features present in patients with PD and other synucleinopathies.

## Discussion

In this paper, we report for the first time an analysis of α-syn pathology in human midbrain organoids (hMOs) derived from a patient with PD carrying an *SNCA* gene triplication. We observed that hMOs carrying this CNV exhibited pathological hallmarks of synucleinopathies and PD that included the presence of oligomeric and phosphorylated α-syn aggregates after 100 days in culture. We further showed that these phosphorylated aggregates were localized in both neuronal and glial populations. Finally, we observed a loss of neurons, including DNs, which coincided with an accumulation of glial cells in the *SNCA* triplication hMOs.

Synucleinopathies are a diverse group of neurodegenerative disorders that share a common pathology, consisting of inclusions containing insoluble α-syn within neurons and glia. Missense mutations and multiplications of the *SNCA* locus have been reported in cases of familial PD, and in particular, CNVs have been identified in families with early-onset, autosomal dominant forms of PD. Interestingly the role of *SNCA* levels has also been investigated in idiopathic sporadic PD, where elevated levels of *SNCA*-mRNA have been reported in midbrain tissue and in the DNs of the SN, suggesting a general role for increased *SNCA* levels in PD ^9, 15^. Consistent with this idea, the role of elevated *SNCA* levels were demonstrated in other synucleinopathies. For instance, analysis of *SNCA* mRNA expression in the human temporal cortex in patients with PD and DLB revealed a correlation between the number of α-syn-immunoreactive LBs and the abundance of *SNCA* mRNA. Additionally, it was shown that oligodendrocytes isolated from the brains of patients with MSA expressed elevated levels of *SNCA* mRNA ^53, 54^. Pathophysiological features of PD have also been observed in hMOs derived from patients with mutations in other PD genes. For instance, Smits et al. ^49^ generated hMOs from patients with leucine-rich repeat kinase 2 (LRRK2) G2019S mutations and observed a reduced number and complexity of midbrain DN compared to hMOs derived from healthy controls. Furthermore, Kim and colleagues ^55^ examined the impact of the LRRK2-G209S mutation on pS129Syn levels in hMOs. These findings, in conjunction with the results we presented in this paper, help validate hMOs as a relevant model to study PD and synucleinopathies. However, further work is required to test the influence of other α-syn mutations (A53T, A30P, etc.) on the development of synuclein pathology. For instance, the A53T mutation has been proposed to accelerate α-syn oligomer and aggregate formation ^56^ and therefore may also lead to an earlier or more severe manifestation of synucleinopathy in hMOs.

Complementary to the studies of synucleopathies in the context of other PD gene mutations, additional work is required to further characterize the features of the synucleopathy. In the SN of a patient with DLB, we used Syn-PLA staining to show that oligomeric α-syn colocalized with total α-syn and pS129Syn, in the LBs and LNs structures. However, in hMOs this observation was less pronounced, most likely as at 100 days, hMOs do not form characteristic LB-like structures. As a result, conducting Syn-PLAs on older cultures will be important to determine whether LBs can develop within hMOs at ages beyond 170 days. The observation that pS129Syn was enriched in 170-day compared to 100-days old hMOs, and that neurons were more significantly reduced at later time point (170 days), also supports the idea that older cultures will lead to more pronounced PD phenotypes. Thus, we hypothesize that aging hMOs for longer time periods, up to 12-18 months, could potentially unveil additional features of synucleinopathies.

To further confirm the pathological similarities between hMOs and patients with PD, using a combination of immunostaining and flow cytometry, we localized phosphorylated α-syn to neuronal and glial populations. While α-syn is abundant in brain, localizing mostly at the presynaptic terminals of neurons, the question of whether it is also expressed in glial cells remains controversial ^57^. Thus, it remains unclear whether our detection of α-syn and pS129Syn in glial cells is the result of endogenous expression or uptake of α-syn released from nearby neurons, which would be consistent with the prion-like spreading and propagation model of synucleinopathies. This latter model is consistent with pathological studies in post-mortem PD brain tissue, in which α-syn positive inclusions have been found in both astrocytes and neurons ^58^. In this scenario, it has been suggested that the α-syn oligomers, which are formed in neurons, are subsequently released and taken up by astrocytes ^58, 59^. It is believed that the normal function of astrocytes is to take up α-syn for the purposes of removal and degradation, and to maintain a healthy environment for neuronal function ^60^. High concentrations of extracellular α-syn have been shown to induce inflammatory and stress responses in astrocytes ^60^, resulting in dysregulation of astrocyte function and eventual apoptosis ^61–63^. Intriguingly, we found a tendency towards a higher CD49f^+^ astrocytic population in *SNCA* Tri hMOs. However, further work on the inflammatory response and gliosis in *SNCA* Tri hMOs is required. It is also important to note that certain cell populations such as microglia, which are derived from mesoderm, are absent from hMOs. Exploring the contribution microglia to ɑ-syn pathology in hMOs, possibly by using co-culture approaches, should be the focus of future work.

Importantly, we observed a selective loss of TH-positive DNs at 100 days in the *SNCA* triplication hMOs, coincidentally with a broader loss of neurons and an increase in glia. In our studies we did not detect differences DN numbers at 50 days but did observe a reduction in the overall number of DNs in the hMOs at later time points. It is interesting to note that we also observed reduced TH expression levels in *SNCA* Tri hMOs at early culture stages (30 days) (**Fig S1G**). *SNCA* Tri hMOs were also smaller in size, suggesting that the *SNCA* triplication impacts early developmental stages in hMOs. Similar observations were reported previously in 2D iPSC-derived neural progenitor lines (2D NPCs) where it was shown that the overexpression of α-syn impaired the differentiation of neuronal progenitor cells ^64^. This could be a result of the neurodevelopmental impact of *SNCA* triplication which has been reported by other groups in iPSC-derived models ^64^.

Overall, our findings validated the hypothesis that hMO models of PD faithfully recapitulate pathological phenotypes seen in patients with PD and other synucleinopathies. They also highlight the potential of hMOs to advance our understanding of the underlying mechanisms involved in PD pathogenesis.

## Acknowledgements

We would like to thank Drs. Rosalind Roberts and Nora Bengoa-Vergniory for kindly sharing their protocol for immunofluorescence combined with Syn-PLA which was optimized for paraffin sections. We also want to acknowledge the MNI Microscopy Core Facility for confocal microscope management and maintenance. We thank the C-BIG repository histology core facility for processing and embedding the paraffin tissue blocks.

---

**Figure S1.**
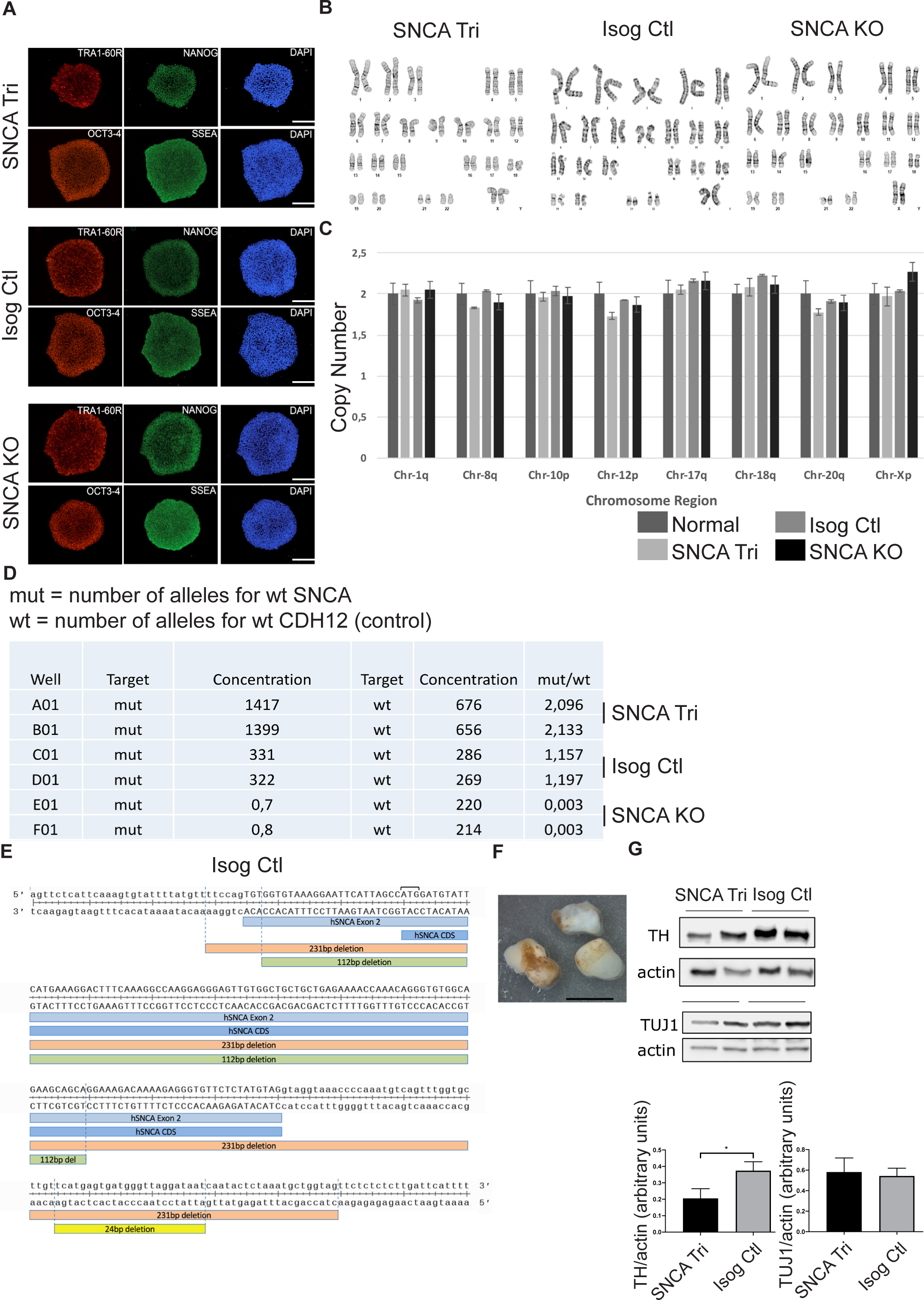
Quality control on *SNCA* iPSCs lines. (**A**) Representative immunostainings for the four key pluripotency markers in iPSC colonies: NANOG, SSEA-4, OCT 3-4, TRA1-60R. Scale bars = 275 μm. (**B**) Karyotyping performed on DNA extracted from iPSC lines showed normal results. (**C**) Genomic stability assay performed on DNA extracted from iPSC lines showed normal copy numbers in commonly mutated sites. (**D**) Digital droplet PCR performed on iPSC DNA samples showed double the number of *SNCA* copies in *SNCA* Tri compared to Isog Ctl, relative to endogenous control *CDH2*. No *SNCA* copies were detected in the *SNCA* KO line. (**E**) Schematic of genetic editing in Isog Ctl line corrected for normal *SNCA* copy number. Exon 2, *SNCA* coding region (CDS) and the deleted regions on each allele are shown (231, 112 and 24 base pairs). Lowercase represents introns and capital letters represent exons. (**F**) Representative image of spontaneous accumulation of neuromelanin granules in *SNCA* Tri 6-month old hMOs. Scale bar = 3 mm. (**G**) Immunoblotting analysis showing presence of dopaminergic marker tyrosine hydroxylase (TH) in 30-day old *SNCA* Tri hMOs and Isog Ctl hMOs, as well as the presence of neuron-specific class III beta-tubulin (TUJ1). All blots were normalized to actin. Quantification by densitometry (arbitrary units) (n=8, mean+/- SEM, Mann Whitney test, two-tailed).

**Figure S2.**
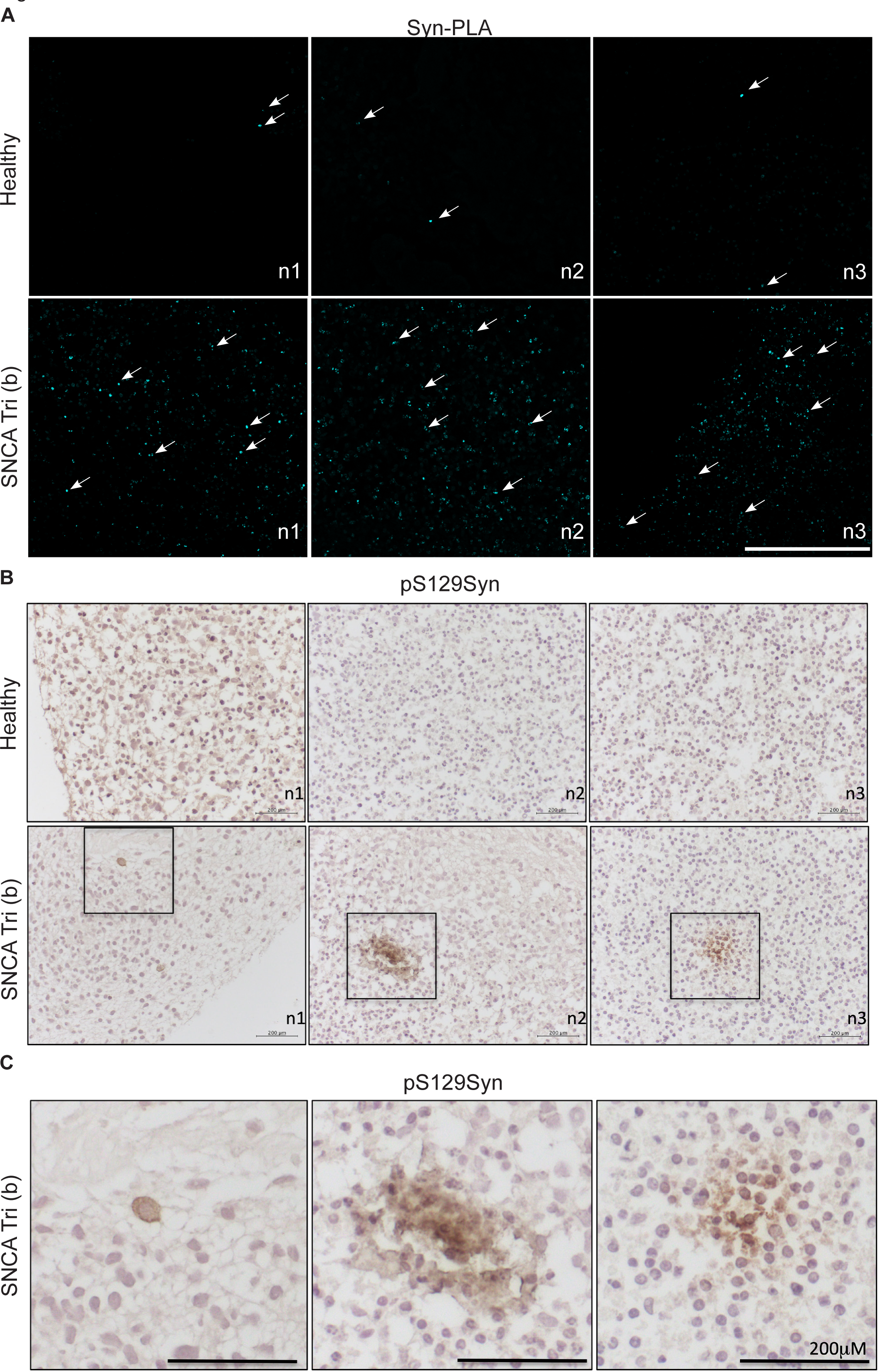
ɑ-syn aggregation in *SNCA* Tri hMOs from another patient (*SNCA* Tri (b)) (**A**) Synuclein proximity ligation assay (Syn-PLA) on paraffin sections showed more α-syn oligomers in 90-day old *SNCA* Tri (b) hMOs (derived from ND34391*H line) compared to healthy individual hMOs (derived from NCRM1 line). Pictures of 3 different organoids for each line (n1, n2, n3) are shown. Scale bar = 600 μm. (**B**) Immunohistochemistry on paraffin sections. Day 90, healthy individual hMOs (derived from NCRM1 line) and *SNCA* Tri (b) hMOs (derived from ND34391*H line) were stained for pS129Syn. Pictures of 3 different organoids for each line (n1, n2, n3) are shown. We observed positive staining in *SNCA* Tri (b) hMOs (squares) but not healthy controls (**C**) Higher magnification of the squares from (*B*) showing pS129Syn staining in 3 different *SNCA* Tri (b) hMOs. Scale bar = 200 μm.

**Figure S3.**
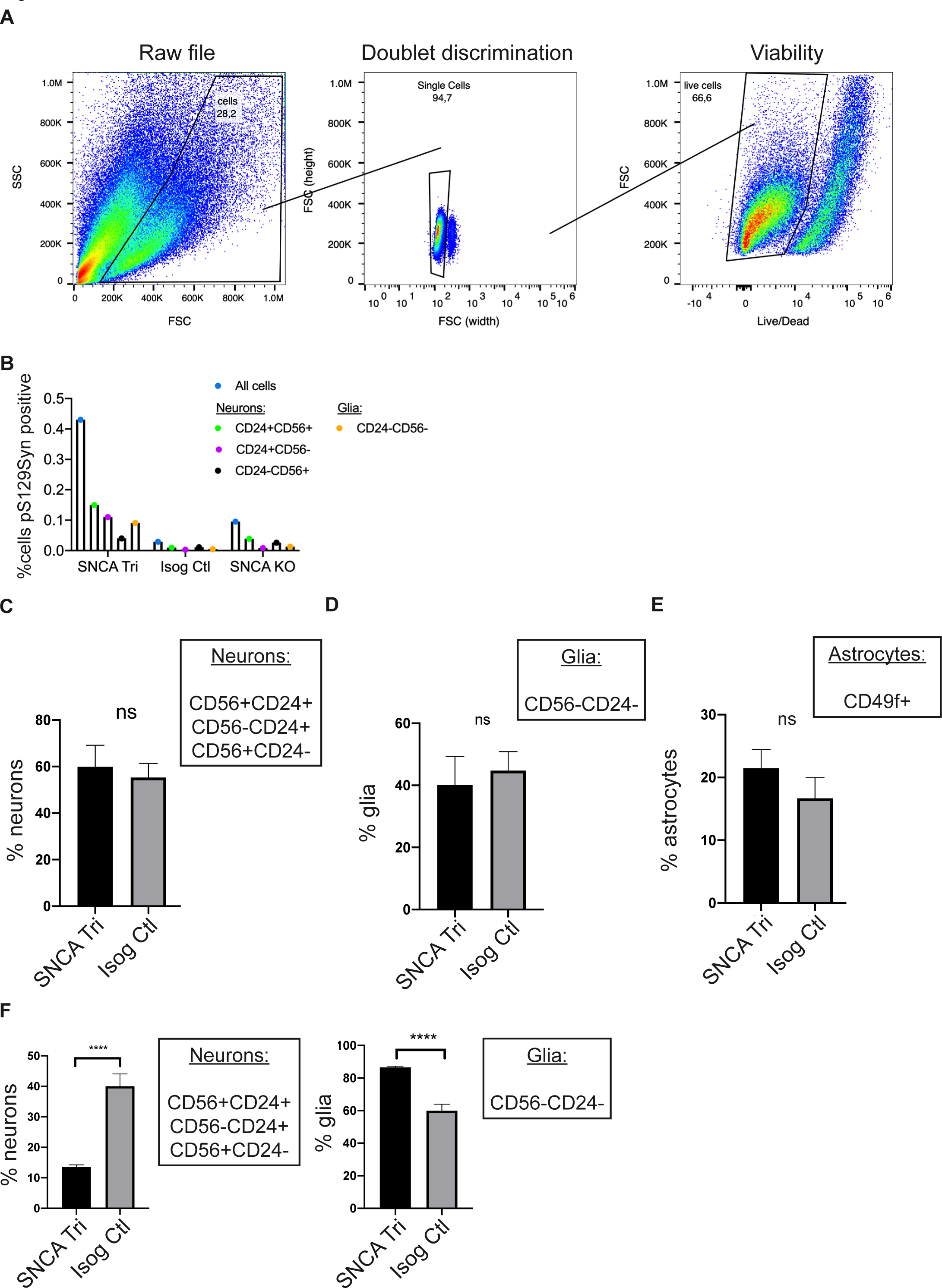
Additional information on flow cytometry. (**A**) Gating strategy for Fig 6D***-F***. (**B**) Proportion of total cells, CD56^+^CD24^+^, CD56^-^ CD24^+^, and CD56^+^CD24^-^neurons, and CD56^-^CD24^-^ glial cells carrying pS129Syn (3 hMOs pooled per line) measured by flow cytometry at 100 days. (**C**) Quantification of total neurons (CD56^+^CD24^+^, CD56^-^CD24^+^, CD56^+^CD24^-^) in 100-day old hMOs (n=3, 3 hMOs pooled per n, 9 organoids total, mean+/-SEM, unpaired t-test, two-tailed). (**D**) Quantification of glia (CD56^-^CD24^-^) in 100-day old hMOs (n=3, 3 hMOs pooled per n, 9 organoids total, mean+/-SEM, unpaired t-test, two-tailed). (**E**) Quantification of astrocytes (CD49f^+^) in 100-day old hMOs (n=3, 3 hMOs pooled per n, 9 organoids total, mean+/-SEM, unpaired t-test, two-tailed). (**F**) Quantification of total neurons (CD56^+^CD24^+^, CD56^-^CD24^+^, CD56^+^CD24^-^) and CD56^-^CD24^-^ glia cells in 170-day old hMOs (n=6, 3 hMOs pooled per n, 18 organoids total, mean+/-SEM, unpaired t-test, two-tailed).

**Table S1.**
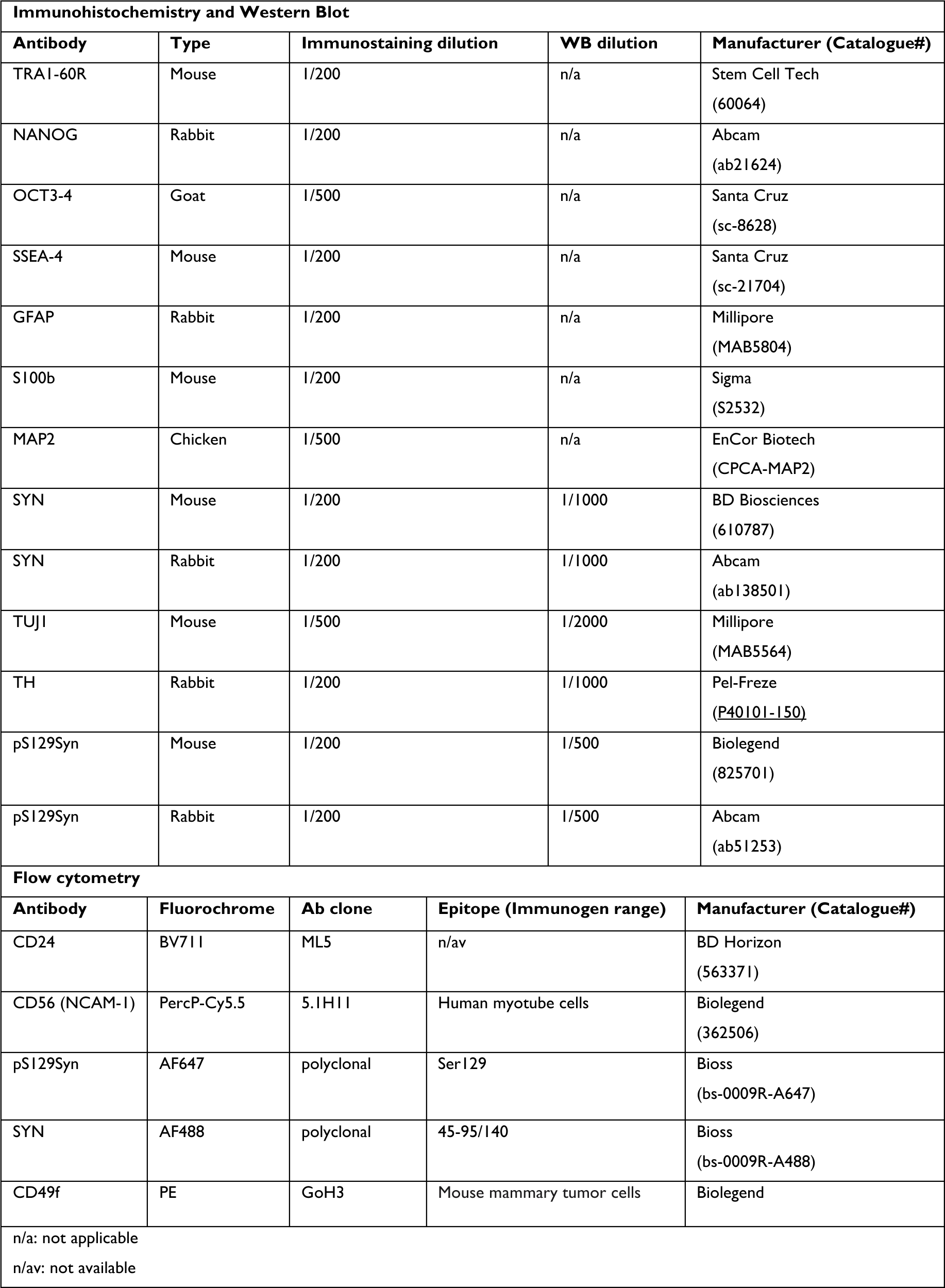
Antibodies.

**Table S2.**
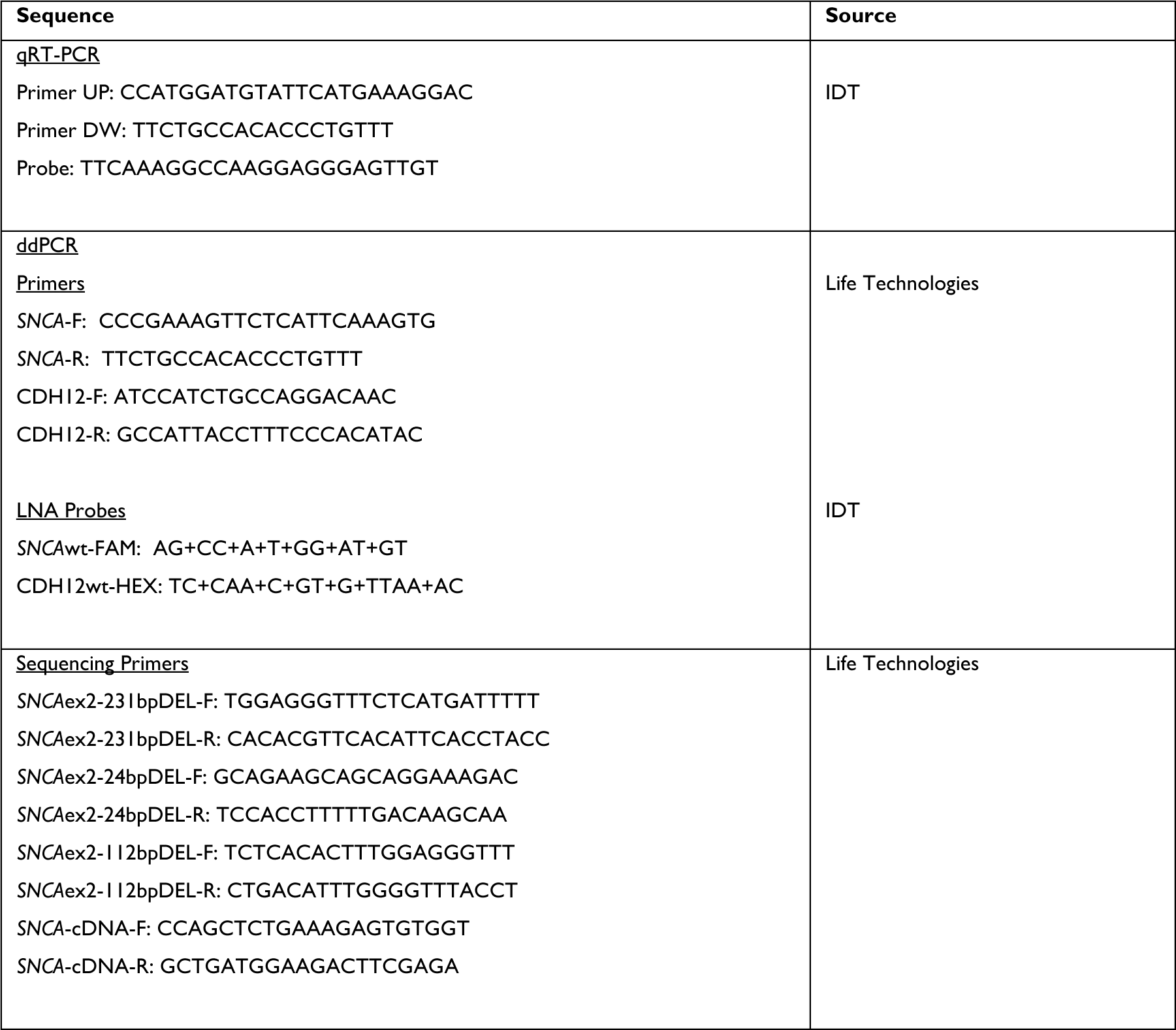
Primers.

## References

1. Lee A, Gilbert RM. Epidemiology of Parkinson’s disease. Neurol Clin. Nov 2016;34(4):955–965. doi:10.1016/j.ncl.2016.06.012

2. Polymeropoulos MH, Lavedan C, Leroy E, et al. Mutation in the alpha-synuclein gene identified in families with Parkinson’s disease. Science. Jun 27 1997;276(5321):2045–7. doi:10.1126/science.276.5321.2045

3. Fujioka S, Ogaki K, Tacik PM, Uitti RJ, Ross OA, Wszolek ZK. Update on novel familial forms of Parkinson’s disease and multiple system atrophy. Parkinsonism Relat Disord. Jan 2014;20 Suppl 1:S29–34. doi:10.1016/S1353-8020(13)70010-5

4. Kasten M, Klein C. The many faces of alpha-synuclein mutations. Mov Disord. Jun 2013;28(6):697–701. doi:10.1002/mds.25499

5. Kruger R, Kuhn W, Muller T, et al. Ala30Pro mutation in the gene encoding alpha-synuclein in Parkinson’s disease. Nat Genet. Feb 1998;18(2):106–8. doi:10.1038/ng0298-106

6. Lesage S, Anheim M, Letournel F, et al. G51D alpha-synuclein mutation causes a novel parkinsonian-pyramidal syndrome. Ann Neurol. Apr 2013;73(4):459–71. doi:10.1002/ana.23894

7. Proukakis C, Dudzik CG, Brier T, et al. A novel alpha-synuclein missense mutation in Parkinson disease. Neurology. Mar 12 2013;80(11):1062–4. doi:10.1212/WNL.0b013e31828727ba

8. Zarranz JJ, Alegre J, Gomez-Esteban JC, et al. The new mutation, E46K, of alpha-synuclein causes Parkinson and Lewy body dementia. Ann Neurol. Feb 2004;55(2):164–73. doi:10.1002/ana.10795

9. Chartier-Harlin MC, Kachergus J, Roumier C, et al. Alpha-synuclein locus duplication as a cause of familial Parkinson’s disease. Lancet. Sep 25-Oct 1 2004;364(9440):1167–9. doi:10.1016/S0140-6736(04)17103-1

10. Singleton AB, Farrer M, Johnson J, et al. Alpha-Synuclein locus triplication causes Parkinson’s disease. Science. Oct 31 2003;302(5646):841. doi:10.1126/science.1090278

11. Farrer M, Kachergus J, Forno L, et al. Comparison of kindreds with parkinsonism and alpha-synuclein genomic multiplications. Ann Neurol. Feb 2004;55(2):174–9. doi:10.1002/ana.10846

12. Nishioka K, Hayashi S, Farrer MJ, et al. Clinical heterogeneity of alpha-synuclein gene duplication in Parkinson’s disease. Ann Neurol. Feb 2006;59(2):298–309. doi:10.1002/ana.20753

13. Ibanez P, Bonnet AM, Debarges B, et al. Causal relation between alpha-synuclein gene duplication and familial Parkinson’s disease. Lancet. Sep 25-Oct 1 2004;364(9440):1169–71. doi:10.1016/S0140-6736(04)17104-3

14. Muenter MD, Forno LS, Hornykiewicz O, et al. Hereditary form of parkinsonism--dementia. Ann Neurol. Jun 1998;43(6):768–81. doi:10.1002/ana.410430612

15. Book A, Guella I, Candido T, et al. A meta-analysis of alpha-synuclein multiplication in familial parkinsonism. Front Neurol. 2018;9:1021. doi:10.3389/fneur.2018.01021

16. Oueslati A. Implication of alpha-synuclein phosphorylation at s129 in synucleinopathies: what have we learned in the last decade? J Parkinsons Dis. 2016;6(1):39–51. doi:10.3233/JPD-160779

17. Lashuel HA, Overk CR, Oueslati A, Masliah E. The many faces of alpha-synuclein: from structure and toxicity to therapeutic target. Nat Rev Neurosci. Jan 2013;14(1):38–48. doi:10.1038/nrn3406

18. Zhang G, Xia Y, Wan F, et al. New perspectives on roles of alpha-synuclein in Parkinson’s disease. Front Aging Neurosci. 2018;10:370. doi:10.3389/fnagi.2018.00370

19. Bengoa-Vergniory N, Roberts RF, Wade-Martins R, Alegre-Abarrategui J. Alpha-synuclein oligomers: a new hope. Acta Neuropathol. Dec 2017;134(6):819–838. doi:10.1007/s00401-017-1755-1

20. Visanji NP, Brotchie JM, Kalia LV, et al. Alpha-synuclein-based animal models of Parkinson’s disease: challenges and opportunities in a new era. Trends Neurosci. Nov 2016;39(11):750–762. doi:10.1016/j.tins.2016.09.003

21. Delenclos M, Burgess JD, Lamprokostopoulou A, Outeiro TF, Vekrellis K, McLean PJ. Cellular models of alpha-synuclein toxicity and aggregation. J Neurochem. Sep 2019;150(5):566–576. doi:10.1111/jnc.14806

22. Agarwal D, Sandor C, Volpato V, et al. A single-cell atlas of the human substantia nigra reveals cell-specific pathways associated with neurological disorders. Nat Commun. Aug 21 2020;11(1):4183. doi:10.1038/s41467-020-17876-0

23. Smits LM, Magni S, Kinugawa K, et al. Single-cell transcriptomics reveals multiple neuronal cell types in human midbrain-specific organoids. Cell Tissue Res. Jul 31 2020;doi:10.1007/s00441-020-03249-y

24. Webber DACSVVTCJM-SRBJA-ARW-MC. A human single-cell atlas of the substantia nigra reveals novel cell-specific pathways associated with the genetic risk of Parkinson’s disease and neuropsychiatric disorders. bioRxiv. 2020;doi:https://10.1101/2020.04.29.067587

25. Mead BE, Karp JM. All models are wrong, but some organoids may be useful. Genome Biol. Mar 27 2019;20(1):66. doi:10.1186/s13059-019-1677-4

26. Jo J, Xiao Y, Sun AX, et al. Midbrain-like organoids from human pluripotent stem cells contain functional dopaminergic and neuromelanin-producing neurons. Cell Stem Cell. Aug 4 2016;19(2):248–257. doi:10.1016/j.stem.2016.07.005

27. Monzel AS, Smits LM, Hemmer K, et al. Derivation of human midbrain-specific organoids from neuroepithelial stem cells. Stem Cell Reports. May 9 2017;8(5):1144–1154. doi:10.1016/j.stemcr.2017.03.010

28. Chen Y, Dolt KS, Kriek M, et al. Engineering synucleinopathy-resistant human dopaminergic neurons by CRISPR-mediated deletion of the SNCA gene. Eur J Neurosci. Feb 2019;49(4):510–524. doi:10.1111/ejn.14286

29. Chen CX-Q, Abdian N, Maussion G, et al. Standardized quality control workflow to evaluate the reproducibility and differentiation potential of human iPSCs into neurons. bioRxiv. 2021:2021.01.13.426620. doi:10.1101/2021.01.13.426620

30. Mohamed NV, Mathur M, V. Da Silva R, et al. Generation of human midbrain organoids from induced pluripotent stem cells. MNI Open Research. 2021;doi:10.12688/mniopenres.12816.2

31. Schmittgen TD, Livak KJ. Analyzing real-time PCR data by the comparative C(T) method. Nat Protoc. 2008;3(6):1101–8. doi:10.1038/nprot.2008.73

32. Sasaki A, Arawaka S, Sato H, Kato T. Sensitive western blotting for detection of endogenous Ser129-phosphorylated alpha-synuclein in intracellular and extracellular spaces. Sci Rep. Sep 18 2015;5:14211. doi:10.1038/srep14211

33. Billings PC, Sanzari JK, Kennedy AR, Cengel KA, Seykora JT. Comparative analysis of colorimetric staining in skin using open-source software. Exp Dermatol. Feb 2015;24(2):157–9. doi:10.1111/exd.12594

34. Roberts RF, Bengoa-Vergniory N, Alegre-Abarrategui J. Alpha-synuclein proximity ligation assay (AS-PLA) in brain sections to probe for alpha-synuclein oligomers. Methods Mol Biol. 2019;1948:69–76. doi:10.1007/978-1-4939-9124-2_7

35. Maecker HT, Trotter J. Flow cytometry controls, instrument setup, and the determination of positivity. Cytometry A. Sep 1 2006;69(9):1037–42. doi:10.1002/cyto.a.20333

36. Baker D, Hirst AJ, Gokhale PJ, et al. Detecting genetic mosaicism in cultures of human pluripotent stem cells. Stem Cell Reports. Nov 8 2016;7(5):998–1012. doi:10.1016/j.stemcr.2016.10.003

37. Andrews PW, Ben-David U, Benvenisty N, et al. Assessing the safety of human pluripotent stem cells and their derivatives for clinical applications. Stem Cell Reports. Jul 11 2017;9(1):1–4. doi:10.1016/j.stemcr.2017.05.029

38. Merkle FT, Ghosh S, Kamitaki N, et al. Human pluripotent stem cells recurrently acquire and expand dominant negative P53 mutations. Nature. May 11 2017;545(7653):229–233. doi:10.1038/nature22312

39. Arenas E, Denham M, Villaescusa JC. How to make a midbrain dopaminergic neuron. Development. Jun 1 2015;142(11):1918–36. doi:10.1242/dev.097394

40. Doi D, Samata B, Katsukawa M, et al. Isolation of human induced pluripotent stem cell-derived dopaminergic progenitors by cell sorting for successful transplantation. Stem Cell Reports. Mar 11 2014;2(3):337–50. doi:10.1016/j.stemcr.2014.01.013

41. Jaeger I, Arber C, Risner-Janiczek JR, et al. Temporally controlled modulation of FGF/ERK signaling directs midbrain dopaminergic neural progenitor fate in mouse and human pluripotent stem cells. Development. Oct 2011;138(20):4363–74. doi:10.1242/dev.066746

42. Kirkeby A, Grealish S, Wolf DA, et al. Generation of regionally specified neural progenitors and functional neurons from human embryonic stem cells under defined conditions. Cell Rep. Jun 28 2012;1(6):703–14. doi:10.1016/j.celrep.2012.04.009

43. Kriks S, Shim JW, Piao J, et al. Dopamine neurons derived from human ES cells efficiently engraft in animal models of Parkinson’s disease. Nature. Nov 6 2011;480(7378):547–51. doi:10.1038/nature10648

44. Perrier AL, Tabar V, Barberi T, et al. Derivation of midbrain dopamine neurons from human embryonic stem cells. Proc Natl Acad Sci U S A. Aug 24 2004;101(34):12543–8. doi:10.1073/pnas.0404700101

45. Tofoli FA, Semeano ATS, Oliveira-Giacomelli A, et al. Midbrain Dopaminergic Neurons Differentiated from Human-Induced Pluripotent Stem Cells. Methods Mol Biol. 2019;1919:97–118. doi:10.1007/978-1-4939-9007-8_8

46. Fedorow H, Tribl F, Halliday G, Gerlach M, Riederer P, Double KL. Neuromelanin in human dopamine neurons: comparison with peripheral melanins and relevance to Parkinson’s disease. Prog Neurobiol. Feb 2005;75(2):109–24. doi:10.1016/j.pneurobio.2005.02.001

47. Alam P, Bousset L, Melki R, Otzen DE. Alpha-synuclein oligomers and fibrils: a spectrum of species, a spectrum of toxicities. J Neurochem. Sep 2019;150(5):522–534. doi:10.1111/jnc.14808

48. Danzer KM, Krebs SK, Wolff M, Birk G, Hengerer B. Seeding induced by alpha-synuclein oligomers provides evidence for spreading of alpha-synuclein pathology. J Neurochem. Oct 2009;111(1):192–203. doi:10.1111/j.1471-4159.2009.06324.x

49. Smits LM, Reinhardt L, Reinhardt P, et al. Modeling Parkinson’s disease in midbrain-like organoids. NPJ Parkinsons Dis. 2019;5:5. doi:10.1038/s41531-019-0078-4

50. Pruszak J, Ludwig W, Blak A, Alavian K, Isacson O. CD15, CD24, and CD29 define a surface biomarker code for neural lineage differentiation of stem cells. Stem Cells. Dec 2009;27(12):2928–40. doi:10.1002/stem.211

51. Janssens S, Schotsaert M, Manganaro L, et al. FACS-mediated isolation of neuronal cell populations from virus-infected human embryonic stem cell-derived cerebral organoid cultures. Curr Protoc Stem Cell Biol. Feb 2019;48(1):e65. doi:10.1002/cpsc.65

52. Barbar L, Jain T, Zimmer M, et al. CD49f is a novel marker of functional and reactive human ipsc-derived astrocytes. Neuron. Aug 5 2020;107(3):436–453 e12. doi:10.1016/j.neuron.2020.05.014

53. Tagliafierro L, Chiba-Falek O. Up-regulation of SNCA gene expression: implications to synucleinopathies. Neurogenetics. Jul 2016;17(3):145–57. doi:10.1007/s10048-016-0478-0

54. Marti MJ, Tolosa E, Campdelacreu J. Clinical overview of the synucleinopathies. Mov Disord. Sep 2003;18 Suppl 6:S21–7. doi:10.1002/mds.10559

55. Kim H, Park HJ, Choi H, et al. Modeling G2019S-LRRK2 sporadic Parkinson’s disease in 3D midbrain organoids. Stem Cell Reports. Mar 5 2019;12(3):518–531. doi:10.1016/j.stemcr.2019.01.020

56. Mohite GM, Kumar R, Panigrahi R, et al. Comparison of kinetics, toxicity, oligomer formation, and membrane binding capacity of alpha-synuclein familial mutations at the A53 site, including the newly discovered A53V mutation. Biochemistry. Sep 4 2018;57(35):5183–5187. doi:10.1021/acs.biochem.8b00314

57. Filippini A, Gennarelli M, Russo I. Alpha-synuclein and glia in Parkinson’s disease: a beneficial or a detrimental duet for the endo-lysosomal system? Cell Mol Neurobiol. Mar 2019;39(2):161–168. doi:10.1007/s10571-019-00649-9

58. Braak H, Sastre M, Del Tredici K. Development of alpha-synuclein immunoreactive astrocytes in the forebrain parallels stages of intraneuronal pathology in sporadic Parkinson’s disease. Acta Neuropathol. Sep 2007;114(3):231–41. doi:10.1007/s00401-007-0244-3

59. Lee HJ, Suk JE, Patrick C, et al. Direct transfer of alpha-synuclein from neuron to astroglia causes inflammatory responses in synucleinopathies. J Biol Chem. Mar 19 2010;285(12):9262–72. doi:10.1074/jbc.M109.081125

60. Booth HDE, Hirst WD, Wade-Martins R. The role of astrocyte dysfunction in Parkinson’s disease pathogenesis. Trends Neurosci. Jun 2017;40(6):358–370. doi:10.1016/j.tins.2017.04.001

61. Rannikko EH, Weber SS, Kahle PJ. Exogenous alpha-synuclein induces toll-like receptor 4 dependent inflammatory responses in astrocytes. BMC Neurosci. Sep 7 2015;16:57. doi:10.1186/s12868-015-0192-0

62. Anderson JP, Walker DE, Goldstein JM, et al. Phosphorylation of Ser-129 is the dominant pathological modification of alpha-synuclein in familial and sporadic Lewy body disease. J Biol Chem. Oct 6 2006;281(40):29739–52. doi:10.1074/jbc.M600933200

63. Liu M, Qin L, Wang L, et al. Alpha-synuclein induces apoptosis of astrocytes by causing dysfunction of the endoplasmic reticulum Golgi compartment. Mol Med Rep. Jul 2018;18(1):322–332. doi:10.3892/mmr.2018.9002

64. Oliveira LM, Falomir-Lockhart LJ, Botelho MG, et al. Elevated alpha-synuclein caused by SNCA gene triplication impairs neuronal differentiation and maturation in Parkinson’s patient-derived induced pluripotent stem cells. Cell Death Dis. Nov 26 2015;6:e1994. doi:10.1038/cddis.2015.318

